# High-speed, long-term, 4D *in vivo* lifetime imaging in intact and injured zebrafish and mouse brains by instant FLIM

**DOI:** 10.1101/2020.02.05.936039

**Authors:** Yide Zhang, Ian H. Guldner, Evan L. Nichols, David Benirschke, Cody J. Smith, Siyuan Zhang, Scott S. Howard

## Abstract

Traditional fluorescence microscopy is blind to molecular microenvironment information that is present in fluorescence lifetime, which can be measured by fluorescence lifetime imaging microscopy (FLIM). However, existing FLIM techniques are typically slow to acquire and process lifetime images, difficult to implement, and expensive. Here, we present instant FLIM, an analog signal processing method that allows real-time streaming of fluorescence intensity, lifetime, and phasor imaging data through simultaneous image acquisition and instantaneous data processing. Instant FLIM can be easily implemented by upgrading an existing two-photon microscope using cost-effective components and our open-source software. We further improve the functionality, penetration depth, and resolution of instant FLIM using phasor segmentation, adaptive optics, and super-resolution techniques. We demonstrate through-skull intravital 3D FLIM of mouse brains to depths of 300 μm and present the first *in vivo* 4D FLIM of microglial dynamics in intact and injured zebrafish and mouse brains up to 12 hours.

## Introduction

Imaging molecular contrast is essential to continued advances in cellular biology. Fluorescence microscopy has been a significant tool over the past decades in imaging cellular and sub-cellular molecular contrast (*1*). Traditional fluorescence microscopy typically obtains molecular contrast by labeling parts of cells with different fluorophores and imaging emission intensity. However, fluorescence emission contains a wealth of information on the molecular microenvironment that is not captured by the emission intensity but is present in the emission decay lifetime. By performing fluorescence lifetime imaging microscopy (FLIM), fluorophores with overlapping emission spectra can be differentiated, and physiological parameters such as pH, refractive index, ion concentration, dissolved gas concentration, and fluorescence resonance energy transfer (FRET) can be measured (*2, 3*). When combined with multiphoton microscopy (MPM) (*4, 5*), FLIM can provide lifetime measurements *in vivo* with high resolution, deep penetration, and reduced photodamage (*6, 7*). Despite the biologically-relevant information provided by fluorescence lifetime, the widespread use of FLIM in biomedical imaging has been limited due to its slow image acquisition and processing speed, low signal-to-noise ratio (SNR), difficult implementation, and high instrumentation cost (*8–13*).

State-of-the-art time-domain (TD) (*12–16*) and frequency-domain (FD) (*9, 10, 17–19*) FLIM techniques have been developed to overcome the limitations above. The most widely used and commercialized TD-FLIM technique is time-correlated single-photon counting (TCSPC), which extracts the lifetime information from a histogram acquired by repetitively recording the arrival time of each photon (*14–16*). Whereas state-of-the-art TCSPC systems can acquire fluorescence decay data at around 10 μs/pixel, the processing for lifetime information normally takes a much longer time through computationally extensive procedures such as curve fitting and deconvolution. To reduce the data processing time, Ryu et al. employed the analog mean-delay (AMD) method to extract the TD lifetime information through numerical integration and realized real-time visualization of fixed tissues (*12*). Besides TCSPC, Bower et al. utilized a high-frequency digitizer to directly sample the TD fluorescence signal and achieved high-speed imaging of metabolic dynamics in living cells (*13*). While both the AMD and the direct sampling methods reduced the pixel dwell time to below 10 μs, they required expensive high-frequency digitizers (>$10,000), and the limited computer memory and bandwidth prevented them from streaming lifetime images without interruption due to the large amount of data sampled by the digitizers. On the other hand, FD-FLIM techniques are less commonly used or commercialized compared to TD-FLIM approaches due to their relatively difficult implementation and low SNR in lifetime measurements (*8, 11*). Conventional FD-FLIM techniques typically require external modulation sources (usually sinusoidal modulation), high-voltage but imperfect (less than 100% modulation degree) detector gain modulation, sequential phase shifting, specialized electronics, and can achieve lifetime image acquisition and processing as fast as 40 μs/pixel (*9, 10, 17, 18*). Recently, Raspe et al. demonstrated siFLIM, an FD-FLIM technique capable of video-rate (around 10 frames per second) lifetime imaging of living cells (*19*). However, siFLIM requires a dedicated camera, and it relies on a widefield microscope with no optical sectioning capabilities; therefore, it cannot be used to acquire 3D lifetime images. Overall, due to the limitations in acquisition or processing speed, SNR, or optical sectioning capabilities, so far, only a few existing FLIM systems have demonstrated 3D lifetime imaging; no systems, however, have demonstrated long-term, time-lapse 3D (i.e., 4D) *in vivo* FLIM through-skull deep in the brain, where useful ballistic photons are scarce due to severe scattering, and prolonged imaging time is not feasible because of animal movements (*20*).

Here, we demonstrate instant FLIM, a novel FD-FLIM technique that utilizes analog signal processing to simultaneously address the challenges in image acquisition and processing, SNR, implementation, and cost faced by existing FLIM methods. We present instant FLIM as a tool that provides real-time streaming of two-photon fluorescence intensity, lifetime, and phasor imaging data. The word “instant” refers to acquiring FLIM data simultaneously with instantaneous processing, where lifetime images and phasor plots (*3, 21*) are instantaneously generated without recording the fluorescence decay curves, thus eliminating the limitations not only in the speed but also the computer memory and bandwidth faced by state-of-the-art FLIM techniques (*12, 13*). We show that an instant FLIM system can be easily implemented as an upgrade to an existing two-photon laser scanning microscope for under $2,500 using cost-effective off-the-shelf components and our open-source, highly modularized, and user-friendly software packages. We also show that the imaging functionality, penetration depth, and resolution of an instant FLIM system can be further improved using phasor segmentation (*22*), adaptive optics (*23*), and super-resolution FLIM (*24*) techniques. Finally, employing an instant FLIM system, we demonstrate intravital 3D lifetime imaging of mouse brains through-skull to depths of 300 μm, and present the first long-term, *in vivo* 4D lifetime imaging of microglial dynamics in intact and injured larval zebrafish and adult mouse brains up to 12 hours.

## Results

### Instant FLIM system and principle

Built upon on our preliminary work presented in (*25*), instant FLIM uses a radio frequency (RF) analog signal processing approach, where the two-photon excitation fluorescence (2PEF) signal is split to four ways and mixed with the phase-shifted 80 MHz reference signals from the Ti:sapphire laser in a multiplexing manner (Fig. 1A; see Fig. S1 and Fig. S2 for details). It is essentially a homodyne FD-FLIM method to extract the lifetime information from the first harmonic of the 80 MHz laser repetition in fluorescence. We employed the reference signal from the laser because the laser repetition rate had a ±1 MHz variance so a fixed-frequency external reference signal could introduce errors, and the implementation was cost-effective. In instant FLIM, the four paths of the fluorescence and reference signals are operated independently and simultaneously during each measurement, and the mixers’ outputs are DC signals that can be digitized by almost any data acquisition devices. The parallel, analog signal processing enabled simultaneous acquisition and instantaneous processing of 2PEF intensity and lifetime images and phasor plots for single- or multi-exponential decay analysis (Fig. 1B; see Section S1 and Section S2 for details). Consequently, no additional acquisition time compared to conventional two-photon intensity measurement was required in instant FLIM, and it was deconvolution-free and fit-free, as lifetime images and phasor plots can be generated instantaneously through basic matrix operations with no extra computation time or memory, eliminating the bottleneck of state-of-the-art FLIM systems (*12, 13*).

**Fig. 1.**
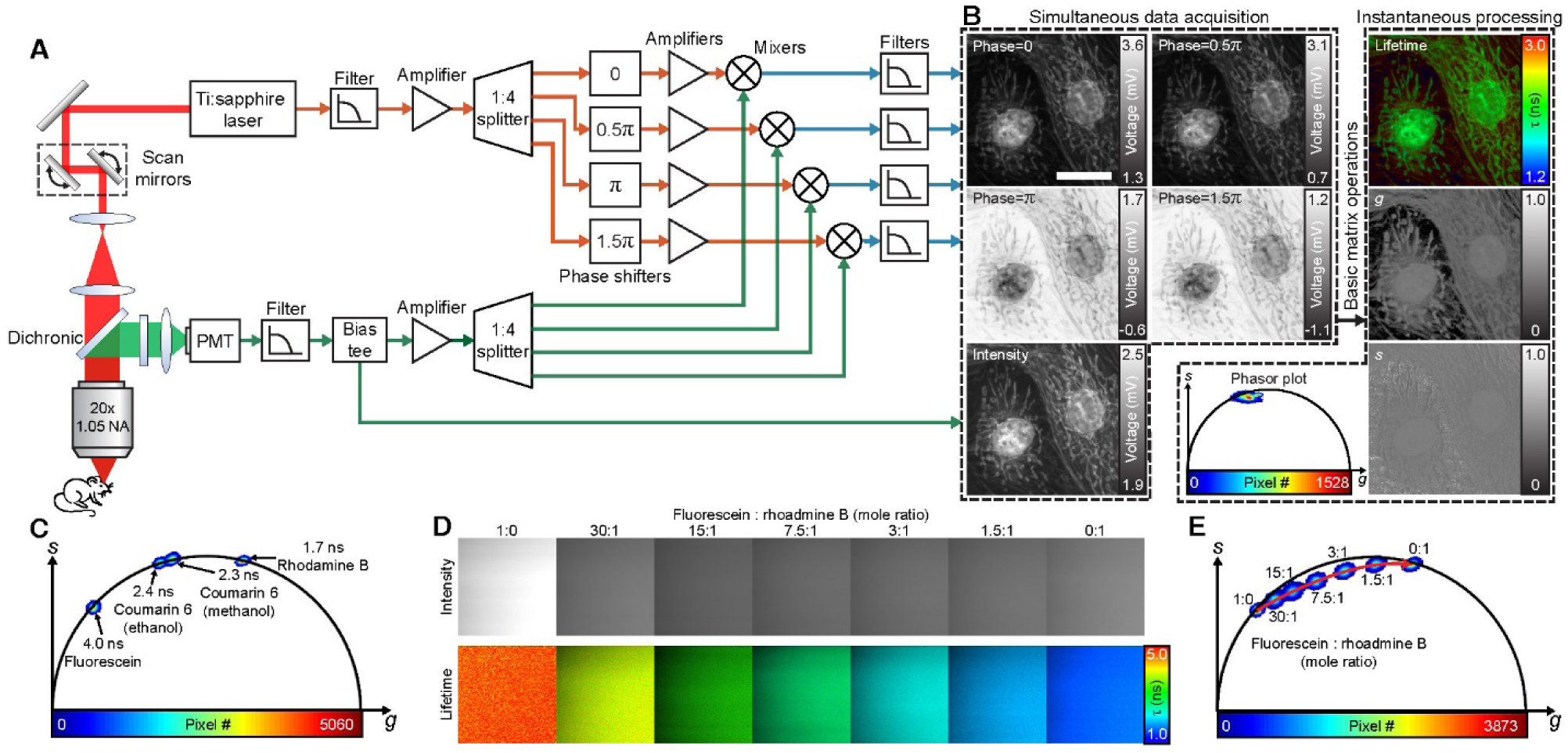
Instant FLIM system. (**A**) Brief diagram of an instant FLIM system. Excitation (red) is generated by a femtosecond laser and sent to a custom-built two-photon laser scanning microscope. Two-photon excitation fluorescence (green) is collected by a photomultiplier tube (PMT), whose signal is filtered and separated to two parts by a bias tee; the RF part is amplified, split to four ways, and mixed with the split laser reference signals with different phases introduced by four phase shifters; the filtered mixer signals and the bias tee’s DC part are then simultaneously digitized by a data acquisition card. (**B**) Left, raw analog signals acquired simultaneously while imaging fixed BPAE cells. Right, lifetime, *g*, and *s* images and phasor plots generated instantaneously through basic matrix operations. (**C**) Phasor plot of four fluorescence lifetime standards. (**D**) Intensity and lifetime images of fluorophore solutions mixed with different mole ratios of fluorescein and rhodamine B. (**E**) Phasor plot of the fluorophore mixtures in (D). The red arrow indicates how the phasors shift when more rhodamine B solution is added to the mixture. Scale bar, 20 μm.

Fluorescence lifetime images alone are usually not enough to resolve the heterogeneity of fluorophores with multi-exponential decays, as different fluorophore compositions could result in the same lifetime measurements. Phasor plots, on the other hand, can resolve the fluorophore heterogeneity because different fluorophore compositions can alter the phasor components (horizontal coordinates *g* and vertical coordinates *s*) even if the average lifetime might be unaltered (*21*). Conventionally, phasor plots are acquired by performing Fourier transforms on fluorescence decay signals. Instant FLIM enables phasor plots generation at the same speed as intensity images through simultaneous data acquisition and instantaneous signal processing without computing Fourier transforms. To confirm the accuracy of phasor measurements of the instant FLIM system, we acquired the phasor plots of four fluorophores with single-exponential decays, which are usually used as fluorescence lifetime standards as their lifetime values are stable and well-studied (*26, 27*). As shown in Fig. 1C, the phasors of these single-exponential fluorophores all located on the universal semicircle, and their lifetime measurements matched the expected values. We then used the instant FLIM system to acquire the intensity and lifetime images and phasor plots of heterogeneous fluorophore mixtures consisting of fluorescein and rhodamine B: whereas the lifetimes of the mixtures of different mole ratios of fluorescein and rhodamine B, e.g., 15:1 and 7.5:1, were hard to distinguish (Fig. 1D), their difference in the phasor plot (Fig. 1E) was evident and could be used to differentiate the two mixtures.

We next analyzed the SNR performance in lifetime measurements of an instant FLIM system and compared it with conventional FD-FLIM techniques (see Section S3 for details). The SNR performance was quantified using the *F*-value, i.e., photon economy, a widely used figure of merit to describe the sensitivity of FLIM techniques. The *F*-value is defined as the ratio of the uncertainties in lifetime and intensity measurements: a smaller *F*-value is desired as it represents a more accurate lifetime measurement with better SNR performance; there is, however, a lower limit of 1 on an *F*-value, which only exists in an ideal shot-noise-limited FLIM system where the available photons are utilized as efficiently as physically possible (*8*). Through Monte Carlo simulations and analytical error-propagation analyses, we showed that instant FLIM has a lower *F*-value, and therefore superior SNR performance, when compared to conventional FD-FLIM techniques for all fluorophores with a lifetime shorter than 4.5 ns. In fact, the *F*-value of instant FLIM approaches the ideal limit of 1, which equals to that of TCSPC. The superior SNR performance of instant FLIM was mainly achieved by efficiently utilizing the intrinsic 80 MHz mode-locked femtosecond laser pulses as the modulation source and employing the low-power external RF mixing, instead of the detector gain modulation used in conventional FD-FLIM systems (*8, 11*).

With the simultaneous image acquisition, instantaneous data processing, and superior SNR performance, in this work, we demonstrated acquisition and processing of 2PEF intensity, lifetime, and phasor imaging data altogether in 12 μs/pixel using a pair of galvo scanners (Thorlabs GVS002) (Table S1). The 12 μs/pixel acquisition time in our instant FLIM system was mainly limited by the bandwidth of the galvo scanners, while the data processing time was around 2 ns/pixel running on an Intel Core i7-8750H 2.20 GHz laptop PC, which was orders of magnitude shorter than the image acquisition time. The pixel dwell time could be greatly reduced if resonant scanning mirrors were used to replace the galvo scanners (*13*), where the intensity of the excitation laser beam needed to be increased accordingly to compensate for the signal loss. We used galvo scanners, instead of resonant mirrors, mainly because galvo scanners are the most widely used scanners in conventional two-photon laser scanning microscopes. Nevertheless, thanks to the modular design of the cost-effective hardware (Fig. S1) and our open-source software (*Instant-FLIM-Control* and *Instant-FLIM-Analysis*) (Fig. S5), modifications to an instant FLIM system such as replacing the galvo scanners with resonant mirrors can be easily implemented.

### Instant FLIM with improved functionality, depth, and resolution

To make instant FLIM a powerful tool for 4D *in vivo* lifetime imaging in deep brains, we further improved its imaging functionality, depth, and resolution using phasor-based image segmentation, adaptive optics (AO), and super-resolution FLIM techniques, respectively. Phasor-based image segmentation techniques utilized phasor plots to segment pixels with similar phasor components (*g* and *s*) in a raw image to resolve fluorescence heterogeneity (*3, 21*), thus improving the functionality of instant FLIM as an imaging tool. Here, we demonstrated two complementary approaches to implement phasor-based image segmentation: the manual phasor labeling (*21*) and the automatic phasor clustering techniques (*22*) (Fig. S6, A and B). In the phasor labeling approach, regions of interest (ROIs) were manually drawn on the phasor plot and the corresponding pixels in the raw image were labeled with different colors (*21*). Fig. 2A shows an example of how different cellular structures in a 3D instant FLIM stack of a fixed Cx3cr1-GFP/+ mouse brain could be segmented into different pseudo-colored groups by manually labeling their phasors with ROIs on the phasor plot in Fig. 2B (Movie S1). On the other hand, the phasor clustering approach segmented an instant FLIM image by applying an unsupervised machine learning approach, i.e., K-means clustering, to automatically cluster the phasors into a specified number of groups (e.g., K=3 as shown in Fig. S6B) and labeling the corresponding pixels in the raw image with different colors (*22*). The two segmentation methods are complementary: the phasor labeling approach is flexible but could lead to biased results, while the phasor clustering approach is unbiased and yet choosing a proper K value might require multiple trials.

**Fig. 2.**
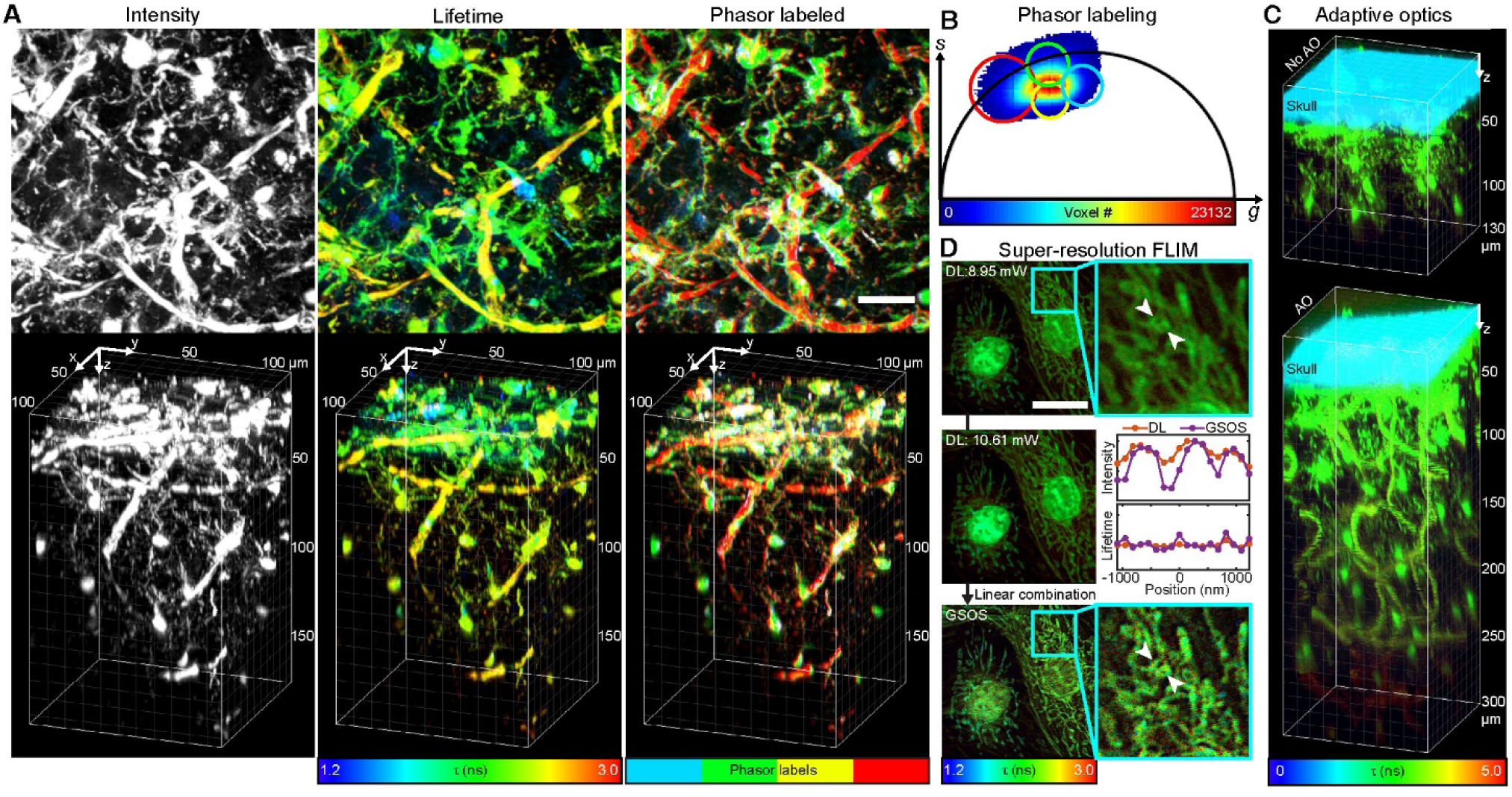
Instant FLIM with phasor labeling, adaptive optics, and super-resolution techniques. (**A**) Maximized z-projections (top) and 3D reconstructions (bottom) of the 2PEF intensity, lifetime, and phasor labeled stacks of a fixed Cx3cr1-GFP/+ mouse brain acquired simultaneously with an instant FLIM system. (**B**) Demonstration of the phasor labeling technique, where ROIs are drawn on the phasor plot of the fixed mouse brain to label different cellular structures in the phasor labeled stack in (A). (**C**) 3D reconstructed lifetime stacks of the intact brain in a living Cx3cr1-GFP/+ mouse acquired with an instant FLIM system without (top) and with (bottom) the adaptive optics (AO) technique. (**D**) Demonstration of super-resolution FLIM by applying the generalized stepwise optical saturation (GSOS) technique in an instant FLIM system, where the super-resolution GSOS lifetime image of fixed BPAE cells is obtained by linear combining two diffraction-limited (DL) instant FLIM images acquired with different excitation powers. Insets, magnified views. Middle, line profiles of the normalized intensity and lifetime values at the positions of the white arrowheads in the DL and GSOS images. Scale bars, 20 μm.

We also added a sensorless AO setup as an optional module to our instant FLIM system (Fig. S1) to improve the penetration depth of our imaging system, which could be used to compensate for the optical aberrations in *in vivo* through-skull imaging deep in mouse brains (*23*). We used one of three optimization algorithms (see Section S4 for details) to iteratively adjust the parameters of the wavefront shaping element, i.e., a deformable mirror in our setup, to improve the image quality quantified by an optimization metric (*28, 29*). We acquired a 3D lifetime stack of the brain, through-skull, in a living Cx3cr1-GFP/+ mouse using the instant FLIM system combined with the AO setup, and compared it with the 3D lifetime stack acquired from the same mouse but without AO (Movie S2). As shown in Fig. 2C, when there was no AO, our instant FLIM system could only achieve a penetration depth of 130 μm for through-skull 3D lifetime imaging in a living mouse brain; in comparison, by adding a well-optimized AO module into our instant FLIM system, the penetration depth for through-skull 3D FLIM in the same mouse was 300 μm, which was more than twice deeper than the depth without AO.

Furthermore, we demonstrated super-resolution FLIM with our instant FLIM system using generalized stepwise optical saturation (GSOS) microscopy (*24*), which used two instant FLIM images acquired at different excitation powers to generate a super-resolution lifetime image with preserved lifetime information and a 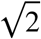-fold increase in spatial resolution (see Section S5 for details). As shown in Fig. 2D, after linear combining two diffraction-limited (DL) lifetime images acquired with excitation powers of 8.95 mW and 10.61 mW, respectively, a super-resolution GSOS lifetime image of the same field-of-view can be generated. Whereas we have utilized GSOS to generate 2D super-resolution FLIM images of fixed cells like the ones in Fig. 2D (*24*), we could not demonstrate GSOS in 3D due to the excessive time required to generate multiple 3D FLIM stacks under different excitation powers using conventional FLIM systems. In instant FLIM, however, due to the analog signal processing, 3D FLIM stacks could be generated at the same speed as 3D intensity stacks. Therefore, by using GSOS in an instant FLIM system, generating a 3D super-resolution FLIM stack only took twice the time of acquiring a 3D two-photon intensity stack. In Fig. S8, we generated a 3D super-resolution FLIM stack by applying GSOS to a pair of DL instant FLIM stacks of a fixed Cx3cr1-GFP/+ mouse brain acquired with excitation powers of 12.31 mW and 13.44 mW, respectively. As shown in Movie S3 and Fig. S8D, the cellular structures in the GSOS stack were better resolved with higher lateral and axial resolutions compared to the ones in the DL stacks.

### *In vivo* 4D instant FLIM imaging in intact zebrafish and mouse brains

We characterized the *in vivo* performance of instant FLIM by measuring the fluorescence lifetime and phasor components of green fluorescence protein (GFP) expressed in microglia in living zebrafish and mouse brains, as microglia are critical for the functionality of the brain in homeostatic and disease states (*30, 31*). The GFP lifetime and the corresponding phasor coordinates are indicators of the intracellular microenvironment and cellular status (e.g., temperature (*32*), refractive index (*33*), stress (*34*), and apoptosis (*35*)). Before measuring GFP lifetimes *in vivo*, we performed an *in vitro* instant FLIM experiment by imaging GFP expressed in living MDA-MB-231-GFP cells under 18 different temperatures from 18.1 °C to 46.7 °C (Fig. S6C). Whereas the intensity images showed no clear distinctions under different temperatures, the fluorescence lifetimes decreased monotonically as temperature increased, consistent with the results reported in (*32*). We also used the phasor labeling (Fig. S6D) and phasor clustering (Fig. S6E) techniques to segment the images and demonstrated that the GFP phasors under different temperatures located differently on the phasor plot. The results confirmed that instant FLIM was able to detect changes in the intracellular microenvironment through lifetime and phasor measurements.

To test instant FLIM as an *in vivo* lifetime imaging modality, we first imaged *Tg(pu1:gfp)* zebrafish at 4 days post fertilization (dpf) using regulatory sequences of *pu1* to express GFP in myeloid cells including microglia and macrophages (Fig. 3A). So far, other than anatomical locations, identifiable markers that distinctly label these cells are limited. To determine if we could detect differences in GFP lifetimes within these cells, we imaged neural regions where both central nervous system (CNS) and peripheral nervous system (PNS) located myeloid cells could be detected and identifiable by their anatomical locations. The instant FLIM measurement reported at least two distinct GFP lifetime profiles using the phasor approach (Fig. 3, B-D). The phasors of PNS-located myeloid cells clustered distinctly compared to that of CNS-located myeloid cells, which also segregated into at least two distinct clusters by their *g* component, suggesting that lifetime could report heterogeneity among myeloid cells. To investigate if lifetime differences were presented in distinct subdomains of individual CNS-located myeloid cells over time, we performed 4D instant FLIM by collecting 3D stacks every 15 minutes for 12 hours (Fig. 3, E and F). The 4D lifetime images in Fig. 3F show that the lifetimes of individual cellular subdomains changed over time as the cells migrated (white arrowheads denote subdomains with increased lifetimes over time). However, these changes largely occurred within the cellular subdomains, whereas the average lifetimes of the cells remained stable during homeostasis (Fig. 3E).

**Fig. 3.**
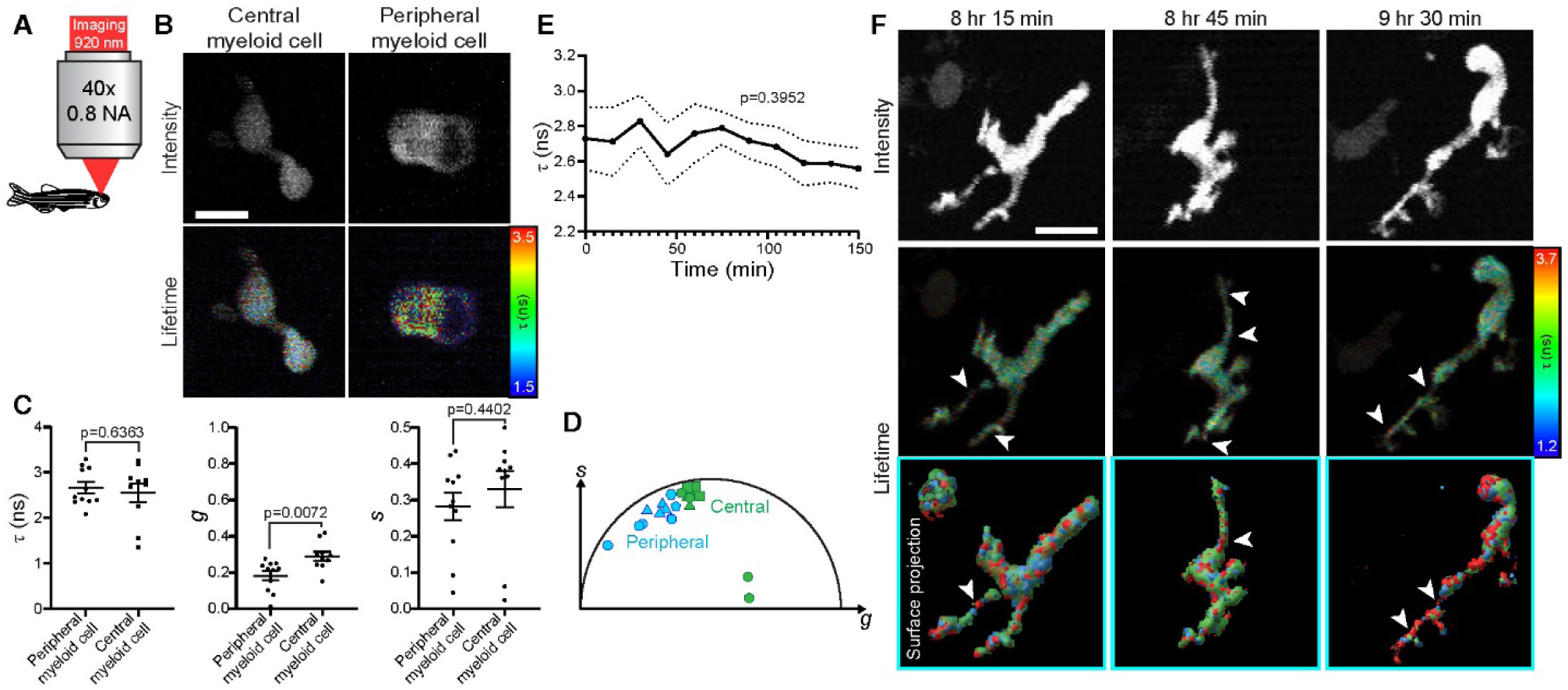
*In vivo* instant FLIM imaging in intact zebrafish brains. (**A**) Larval *Tg(pu1:gfp)* zebrafish at 4 dpf were imaged with a 920 nm laser using instant FLIM. (**B**) 2PEF intensity and lifetime maximized z-projection images of the CNS-located and PNS-located myeloid cells in the zebrafish. (**C**) Graphs of average lifetime (*τ*) and phasor (*g* and *s*) values in CNS-located (n=10 cells) and PNS-located (n=11 cells) myeloid cells. CNS-located cells have a larger *g* phasor component. (**D**) Plot representing the phasor locations of CNS-located (green) and PNS-located (blue) myeloid cells by their *g* and *s* phasor components. Each shape represents a measurement taken from different individual animals. (**E**) Average lifetime values of the zebrafish microglia over 150 minutes (n=8 cells). (**F**) 2PEF intensity and lifetime maximized z-projection images from a 12-hour 4D instant FLIM movie of the zebrafish microglia. Bottom, surface projections representing lifetime subdomains within the cell, where colors denote approximate lifetime values. White arrowheads denote cellular subdomains with increased lifetimes. Scale bars, 10 μm. (C) uses two-sided Student’s t-tests. (E) uses one-way ANOVAs.

We repeated this analysis using skull-thinned Cx3cr1-GFP/+ adult mice whose microglia were labeled with GFP (Fig. 4A). We also retro-orbitally injected Texas Red-Dextran into the animal to distinguish blood vessels from microglia. These adult microglia displayed dynamic and complex branching emanating from the cell body (Fig. 4B). Given that microglia in larval zebrafish displayed distinct lifetime subdomains, we hypothesized that adult microglia could also have distinct lifetimes in their cell bodies and surveilling protrusions. To test this, we measured *τ, g*, and *s* and compared these values between the cell bodies and protrusions. While we did not detect differences in *g* and *s*, the surveilling protrusions displayed increased *τ* compared to that of the cell bodies (Fig. 4,C and D), indicating that the protrusions had distinct changes in their microenvironment while interacting with other neural cell types. We also employed 4D instant FLIM by taking 3D stacks of the mouse microglia every two minutes for six minutes (Fig. 4, E and F). These images demonstrated that while the microglia had dynamic processes, under homeostatic conditions, the lifetimes of the cell bodies and surveilling protrusions were stable.

**Fig. 4.**
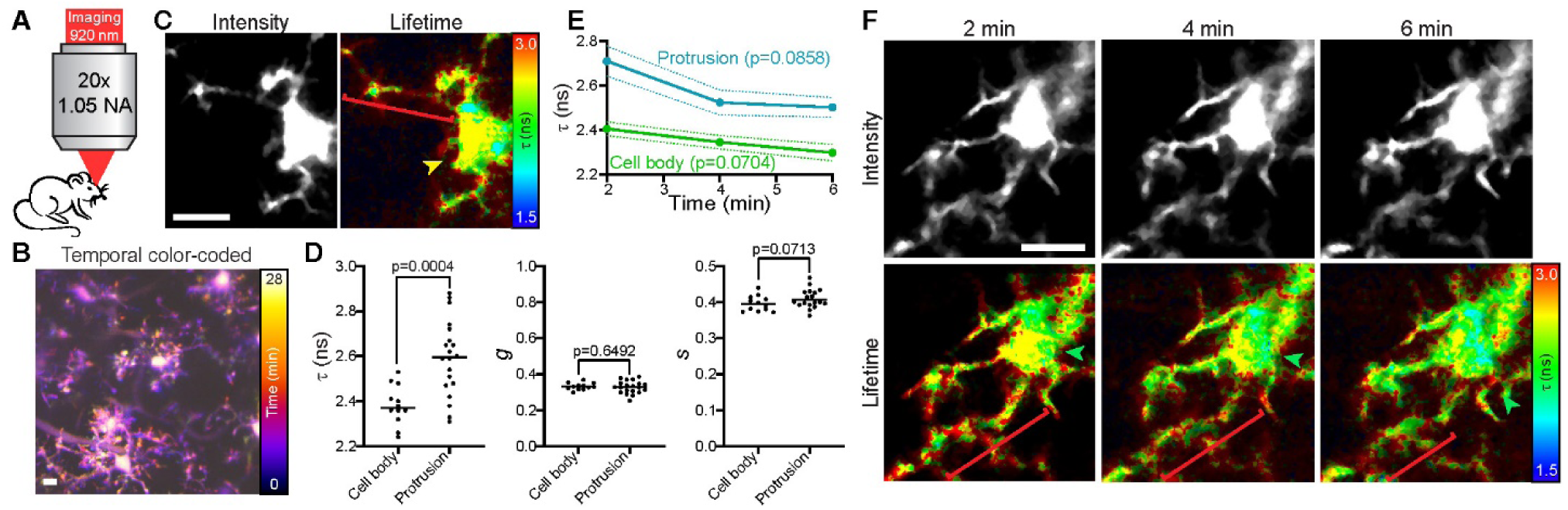
*In vivo* instant FLIM imaging in intact mouse brains. (**A**) Adult Cx3cr1-GFP/+ mice were imaged with a 920 nm laser using instant FLIM. (**B**) 2PEF intensity maximized z-projection image pseudo-colored to represent the temporal dynamics of the microglia. The temporal scale is shown on the right. (**C**) 2PEF intensity and lifetime maximized z-projection images of the mouse microglia. Bracket denotes the microglial protrusion, arrowhead denotes the cell body, and their colors represent approximate lifetime values. (**D**) Graphs of average lifetime (*τ*) and phasor (*g* and *s*) values in microglial cell bodies (n=12) and protrusions (n=20). (**E**) Average lifetime values of microglial cell bodies (green) and protrusions (blue) over 6 minutes (n=3 cells). (**F**) 2PEF intensity and lifetime maximized z-projection images from a 4D instant FLIM movie of the mouse microglia. Brackets denote microglial protrusions, arrowheads denote cell bodies, and their colors represent approximate lifetime values. Scale bars, 10 μm. (D) uses two-sided Student’s t-tests. (E) uses one-way ANOVAs.

### *In vivo* 4D instant FLIM imaging in injured zebrafish and mouse brains

As the brain’s immune cells, microglia sense injury by continuously surveying the parenchyma with highly motile processes and converging to the site of injury to establish a barrier between the healthy and injured brain tissue (*31, 36*). We next used instant FLIM to determine if we could detect lifetime changes in microglia when they responded to brain injury. To do this, we created lesions in the larval *Tg(pu1:gfp)* zebrafish brain with the femtosecond laser and acquired 4D instant FLIM movies of microglial response to the site of injury by collecting 3D stacks every 15 minutes for 12 hours (Fig. 5A). Consistent with the injury response (*36*), microglia migrated to the site of injury (Fig. 5B; Movie S4). We investigated the phasors extracted from the 4D instant FLIM measurement. As shown in Fig. 5C, the phasors of pre-injury microglia clustered together; from 0 to 60 minutes post-injury (mpi), the phasors segregated into two clusters in opposite directions; this clustering continued after 60 mpi with few phasors resembling pre-injury ones. Meanwhile, in plotting average *τ, g*, and *s* values, we detected decreased *s* components in microglia after injury (Fig. 5D). We next analyzed the lifetimes of whole microglia and their individual cellular subdomains in response to injury (Fig. 5, E and F). After injury, both the microglia and subdomains adjusted their lifetimes, as denoted by the change in average lifetime values (Fig. 5E) and the increase of the number of red subdomains (red denoting longer lifetimes) (Fig. 5F), with continuing changes through 75 mpi. These observations demonstrated instant FLIM’s capacity to detect lifetime changes *in vivo* after physiological challenges such as laser injury.

**Fig. 5.**
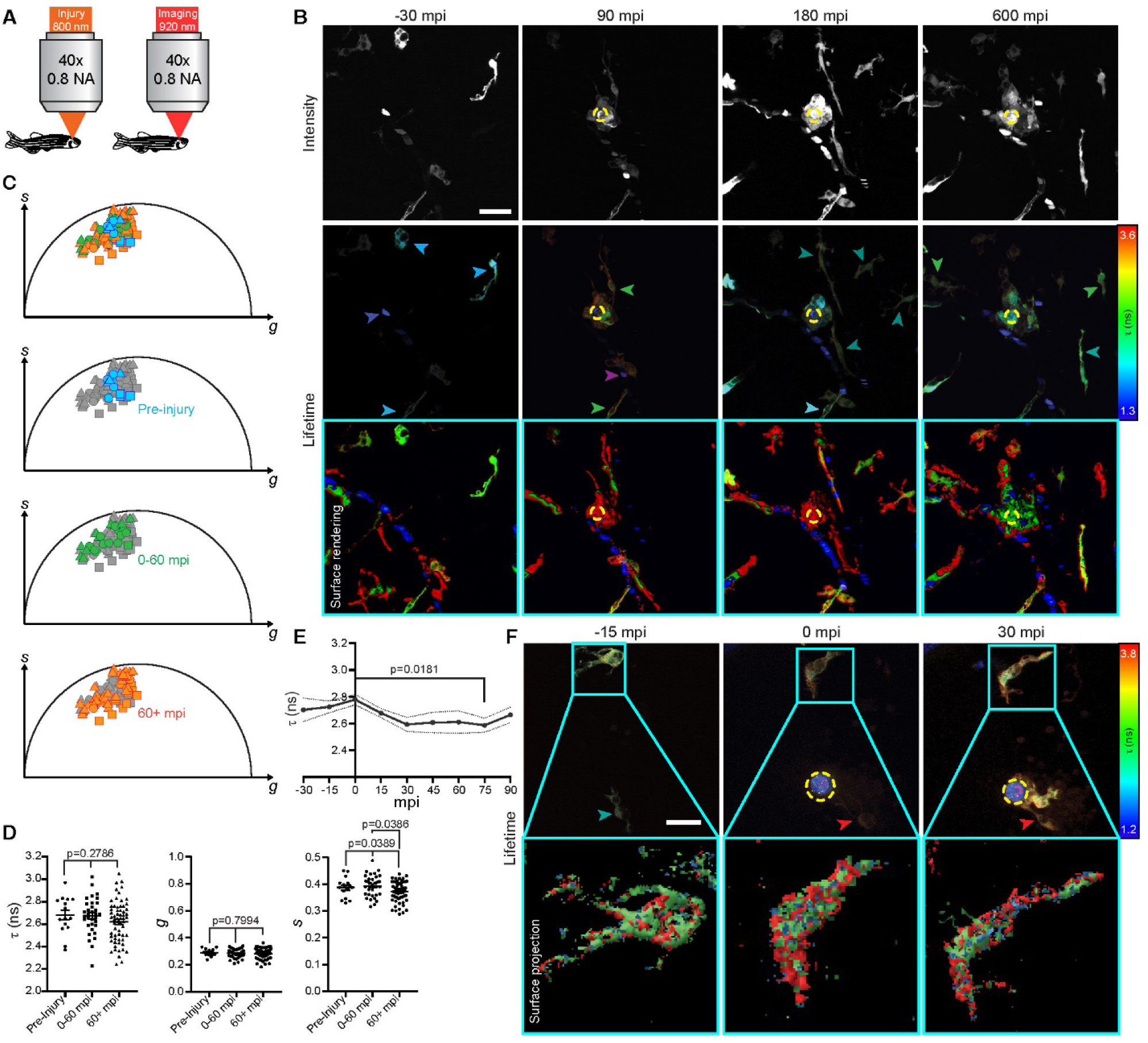
*In vivo* instant FLIM imaging in injured zebrafish brains. (**A**) Larval *Tg(pu1:gfp)* zebrafish at 4 dpf were imaged using instant FLIM following an injury induced by an 800 nm laser. (**B**) 2PEF intensity and lifetime maximized z-projection images from a 12-hour 4D instant FLIM movie in the zebrafish brain before and after laser injury. Mpi, minutes post-injury. Bottom, surface rendering of the lifetime images to better differentiate the cellular structures. Dashed yellow circle denotes the lesion site. Arrowheads denote approximate lifetime values. (**C**) Plot representing the phasor locations of microglia before injury (blue), 0-60 mpi (green), and 60+ mpi (orange). Each shape represents a measurement taken from different individual animals. (**D**) Graphs of average *τ, g*, and *s* values in microglia before injury (n=15 cells), 0-60 mpi (n=33 cells), and 60+ mpi (n=62 cells). (**E**) Average lifetime values of the microglia before and after injury (n=9 cells). (**F**) Lifetime maximized z-projection images from a 4-hour 4D instant FLIM movie of the microglia before and after injury. Insets, surface representation of changing lifetime subdomains within single microglia, where colors denote approximate lifetime values. Dashed yellow circle denotes the lesion site. Arrowheads denote approximate lifetime values. Scale bars, 10 µm. (D) and (E) use Tukey’s HSD tests.

We then used instant FLIM to investigate if similar lifetime changes could be detected in adult mouse microglia when they responded to brain injury. We created lesions in the skull-thinned Cx3cr1-GFP/+ mouse brain with the femtosecond laser and acquired 4D instant FLIM movies of microglial dynamics in response to injury by collecting 3D stacks every two minutes for an hour (Fig. 6A). Adult mouse microglia were markedly more abundant than larval zebrafish microglia. Consistent with the observations reported in (*36*), the laser injury in the adult mouse brain parenchyma induced an instantaneous response from the microglia where processes were extended toward the site of injury (Fig. 6B; Movie S5). This was in contrast with microglia in larval zebrafish which utilized whole-cell migrations to the site of injury. Adult mouse microglial processes surrounded the lesion site similar to the cloaking arrangement of macrophages in the PNS (*37*). We first compared lifetime changes from before the injury (−2 mpi) to after the microglia have extended their processes to the injury site (22 mpi) (Fig. 6C). To do this, we measured the lifetimes and phasor components of the cell bodies, responding protrusions, and protrusion tips which were migrating to the site of injury (Fig. 6D). These measurements did not detect changes in *τ, g*, or *s* from −2 mpi to 22 mpi but did detect differences in *τ* and *g* between each of the cellular subdomains. These data suggested that adult microglia maintained subcellular lifetime and phasor differences while responding to injury (Movie S6). Further, we analyzed lifetime dynamics throughout the process of protrusion extension to cloak the injury site (Fig. 6, E and F). These measurements indicated that lifetimes and phasors in each subdomain of the microglia remained relatively stable during early extension and cloaking of the injury site (Fig. 6E). Considering the stability of lifetime and phasor information of adult mouse microglia in response to injury, we used both the phasor labeling and phasor clustering approaches to segment the 4D instant FLIM voxels into different cellular structures (e.g., cell bodies, protrusions, protrusion tips, etc.) based on the similarity of their phasors and distinguished their responses to the laser injury (Movie S7).

**Fig. 6.**
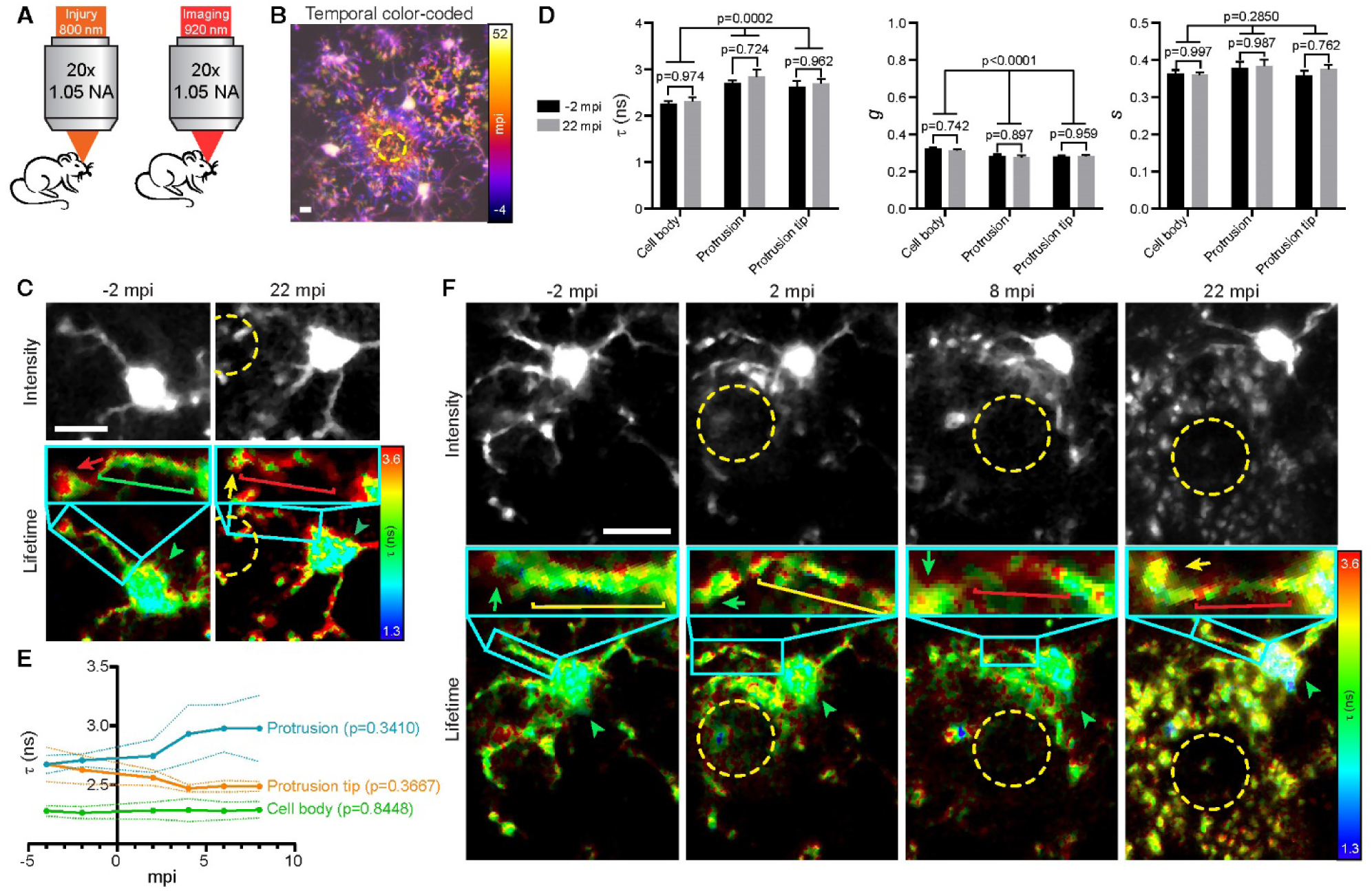
*In vivo* instant FLIM imaging in injured mouse brains. (**A**) Adult Cx3cr1-GFP/+ mice were imaged using instant FLIM following an injury induced by an 800 nm laser. (**B**) 2PEF intensity maximized z-projection image pseudo-colored to represent the temporal dynamics of the microglia in response to injury. The temporal scale is shown on the right. Dashed yellow circle denotes the lesion site. (**C**) Intensity and lifetime maximized z-projection images from a 4D instant FLIM movie in the mouse brain at −2 mpi and 22 mpi. Dashed yellow circle denotes the lesion site. Arrowheads denote cell bodies, brackets denote protrusions, and arrows denote protrusion tips. Colors denote approximate lifetime values. Insets show magnified views of the protrusions. (**D**) Graphs of average *τ, g*, and *s* values in cell bodies (n=5), protrusions (n=5), and protrusion tips (n=5) at −2 mpi and 22 mpi. (**E**) Average lifetime values of cell bodies, protrusions, and protrusion tips before and after injury (n=5 cells). (**F**) Intensity and lifetime maximized z-projection images from a 30-min 4D instant FLIM movie of the microglia before and after injury. Annotations are identical to the ones in (C). Scale bars, 10 µm. (D) uses two-way ANOVAs with multiple comparisons. (E) uses one-way ANOVAs.

Taken together, these results demonstrate the capability of instant FLIM to delineate the dynamics of lifetimes and phasors in diverse cellular subdomains in zebrafish and mouse brains *in vivo*. They also highlight an important contrast in microglial responses to injury early in development and adulthood: microglia in larval zebrafish demonstrated lifetime changes after injury, while this effect was not observed in microglia in adult mice. Whereas the fundamental causes of the GFP lifetime or phasor variations in microglial subdomains are beyond the scope of this article, these variations measured by instant FLIM will be beneficial to distinguish and label cellular subdomains in microglia and understand their roles in the immune surveillance process (*31*).

## Discussion

Through analog signal processing, we have demonstrated instant FLIM as a novel FD-FLIM system that enables simultaneous acquisition and instantaneous processing of 2PEF intensity, lifetime, and phasor imaging data. However, the performance does not come without caveats. First, since instant FLIM utilizes the intrinsic femtosecond laser pulses as the modulation source, the modulation frequency is fixed at 80 MHz; therefore, as shown in Fig. S4C, the SNR performance in lifetime measurements, quantified by the *F*-value, will become worse for fluorophores with longer lifetimes. As a result, instant FLIM will perform best with short-lifetime fluorophores such as rhodamine B and indocyanine green (ICG) (*38*); for long-lifetime fluorophores or samples consisting of a wide range of lifetime components (*39*), the pixel dwell time for instant FLIM measurements should be increased accordingly to compensate for the lower photon economy. Second, while it is advantageous for instant FLIM to generate lifetime images and phasor plots directly without recording fluorescence decay curves to eliminate limitations in computer memory and bandwidth, users who are familiar with TD-FLIM systems may find it less intuitive or not straightforward to interpret the lifetime results. This is a trade-off between functionality and cost, as fluorescence decay curves can only be fully recorded using expensive TCSPC or high-frequency digitizers. Third, due to the electronic and RF components required to implement the analog signal processing in instant FLIM, additional electronic noise such as thermal noise and electromagnetic interference from the environment could be introduced to the lifetime measurement if the instruments were not well connected or shielded. In our experience, however, using low-noise amplifiers and properly shielding the electronics could successfully suppress the additional noise, and shot-noise limited measurements could be achieved.

The four-phase analog signal processing in the current instant FLIM system can be further improved. Theoretically, to extract lifetime information via homodyne FD-FLIM measurements, minimally only three phase images are required, and the lifetime measurement accuracy can be further improved if more phase images, e.g., 12, are employed (*19*). In conventional FD-FLIM systems, these phase images need to be captured sequentially, so acquiring more phase images requires a longer acquisition time. Meanwhile, for more phase images, the processing for lifetime information becomes more complicated and time-consuming, as complex trigonometry calculations or curve fittings are required. In this work, we only acquired four phase images to reduce the cost of the system and the complexity of lifetime and phasor calculations. Nevertheless, following the principle of multiplexing analog signal processing, the current instant FLIM setup could be readily expanded to allow simultaneous acquisition of 12 or more phase images, where the 2PEF signal could be split to 12 or more paths and mixed with the phase-shifted reference signals from the femtosecond laser; therefore, the lifetime measurement accuracy of instant FLIM could be improved while no extra image acquisition time would be needed.

In conclusion, we have presented instant FLIM as a powerful tool for high-speed, long-term, 4D *in vivo* FLIM that allows real-time streaming of 2PEF intensity, lifetime, and phasor imaging data. We have also demonstrated that an instant FLIM system could be combined with phasor labeling and clustering, AO, and GSOS to provide versatile phasor-based image segmentation, deep penetration depths, and super-resolution FLIM performances. In addition, we have demonstrated 3D *in vivo* FLIM of mouse brains through-skull to depths of 300 μm, and we have achieved, to our knowledge, the first long-term, *in vivo*, 4D lifetime imaging of microglia dynamics in intact and injured larval zebrafish and adult mouse brains up to 12 hours. Biologically, the identification of cellular subdomains with different fluorescence lifetime properties of a cytosolic fluorophore was surprising. These domains could be the result of a variety of phenomena such as distinct cell signaling hubs or organelle density. Regardless, these results were accomplished by upgrading a conventional two-photon laser scanning microscope using cost-effective off-the-shelf components with a total cost less than $2,500 and our fully open-source, highly modularized, and user-friendly software packages (*Instant-FLIM-Control* and *Instant-FLIM-Analysis*). As a result, instant FLIM can be easily accessed by many labs and has the potential to enable future discoveries in biology, including the mechanism of lifetime subdomains identified here, in addition to a wide variety of disciplines.

## Materials and Methods

### Experimental setup and analog signal processing

We implemented the instant FLIM system based on a custom-built two-photon laser scanning microscope (Fig. 1A; Fig. S1). The intensity of a mode-locked Ti:sapphire laser (Spectra-Physics Mai Tai BB, 710-990 nm, 100 fs, 80 MHz) is controlled by a neutral density filter (NDF), a motorized half-wave plate, and a polarizing beam splitter (PBS). A power meter is used to measure the excitation power by monitoring a small fraction of the laser beam reflected by a glass slide. A mechanical shutter (Thorlabs SHB1T) is used to block the laser beam when no imaging acquisition takes place. A sensorless adaptive optics (AO) setup consisting of a deformable mirror (Thorlabs DMP40-P01), a beam expander composed of two achromatic doublets (Thorlabs AC254-030-B, AC254-075-B), an achromatic quarter-wave plate (Thorlabs AQWP05M-980), and another PBS is utilized to correct the wavefront of the excitation beam in order to improve the imaging depth (Section S4). The AO setup is optional and can be switched into and out of the optical path using a folding mirror. The laser beam is directed to a conventional two-photon microscopy setup consisting of a pair of galvo scanners (Thorlabs GVS002), a scan lens (Thorlabs AC254-040-B), a tube lens (Thorlabs AC254-200-B), an objective lens (Nikon CFI APO NIR, 40x, 0.8 NA, or Olympus XLPLN25XWMP2, 25x, 1.05 NA), and a motorized stage (Prior OptiScan III). A long pass filter is used to block ambient light from entering the objective. The 2PEF is epi-collected by the objective lens, reflected by a dichroic mirror, filtered through a set of bandpass and short pass filters to eliminate residual excitation, and detected by a photomultiplier tube (PMT) (Hamamatsu H7422PA-40) through a collection lens.

The analog signal processing module in instant FLIM utilizes the fluorescence signal from the PMT and the 80 MHz reference signal from the Ti:sapphire laser to generate the intensity, lifetime, and phasor data simultaneously. As shown in Fig. S2, the current signal from the PMT is amplified and converted to a voltage signal by a transimpedance amplifier (Aricorp DC-100), filtered by a low pass filter (LPF) (Mini-Circuits BLP-90+), and separated into DC and RF parts by a bias tee (Mini-Circuits ZFBT-282-1.5A+). The DC signal (***V***_DC_) is acquired by a data acquisition (DAQ) card (National Instruments PCIe-6323) to reconstruct the intensity image. The RF signal is amplified by a low-noise amplifier (LNA) (Mini-Circuits ZX60-P103LN+) and split to four paths by a 4-way power splitter (Mini-Circuits ZSC-4-3+). Meanwhile, the 80 MHz reference signal from the laser, *v*_ref_ (*t*), is filtered by an LPF to eliminate higher harmonics, amplified by an LNA, and split to four signal paths, *v*_80MHz_(*t*), also by a power splitter. A phase shift is introduced to each reference signal path by a pair of phase shifters (Mini-Circuits JSPHS-150 with TB-152+). Note that each phase shifter can introduce a phase shift within *π* to the 80 MHz signal; connecting them in tandem introduces a total phase shift within 2*π* for each path. The phase shifts are independently controlled by the bias voltages generated by the DAQ card. The four phase-shifted reference signals are then amplified by LNAs and directed to four RF mixers (Mini-Circuits ZAD-3H+). Each mixer takes a reference signal path, *v*_LO_(*t, φ*), to its local oscillator (LO) port and a PMT signal path, *v*_RF_(*t*), to its RF port and generates a mixed signal, *v*_IF_(*t, φ*), on its intermediate frequency (IF) port. The IF signals from each mixer are then filtered to DC, i.e., ***V***_IF_(*φ*), by an LPF (Thorlabs EF502) and measured by another DAQ card (National Instruments PCI-6110). In the end, for each intensity image (2D, 3D, or 4D) acquired, four corresponding mixer images are acquired simultaneously, so the data acquisition is intrinsically synchronized and effect of the PMT timing jitter is minimized. Fluorescence lifetime images as well as phasor plots are then instantaneously generated through basic matrix operations. The DAQ cards are also used to control the motorized half-wave plate, the shutter, and the galvo scanners during the imaging process. The motorized stage and the deformable mirror, on the other hand, are controlled directly by the computer through universal serial bus (USB) ports. Note that all electronic components in the instant FLIM system are impedance matched to 50 Ω to suppress signal reflections, and LNAs are used and the RF components are properly shielded to minimize additional electronic noise introduced to the measurement.

The parts and price list for the implementation of an instant FLIM system is presented in Table S2. The total cost to upgrade an existing two-photon laser scanning microscope (including data acquisition devices) to an instant FLIM system is less than $2,500.

### Simultaneous acquisition and processing of intensity, lifetime, and phasor data

With the analog signal processing in instant FLIM, the fluorescence intensity, lifetime, and phasor data can be acquired simultaneously (Fig. 1B; Section S1). Specifically, the 2PEF intensity image is generated from the DC part of the PMT signal:

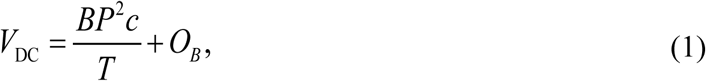

where ***P*** is the excitation power, *c* is the fluorophore concentration, *T* is the modulation period of the Ti:sapphire laser, *B* and ***O***_*B*_ are the conversion loss and offset from the bias tee’s RF&DC to DC ports, respectively. The lifetime and phasor data, on the other hand, are obtained by applying basic mathematical operations on the outputs of the four mixers’ IF ports:

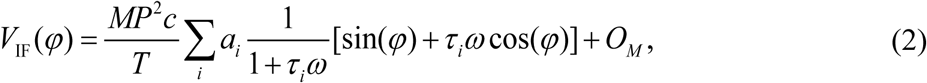

where *φ* is the phase shift introduced by the phase shifters, *a*_*i*_ is the intensity-weighted fractional contribution of the fluorophore with lifetime *τ*_*i*_ (∑_*i*_ *a*_*i*_ = 1), *ω* is the angular modulation frequency, *M* and ***O***_***M***_ are the conversion loss and offset from the mixer’s RF to IF ports, respectively. The average (phase) lifetime (*2*) image is calculated as

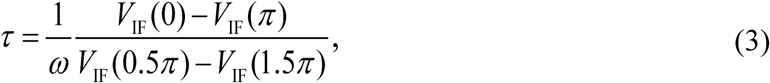

and the components of the phasors are obtained as

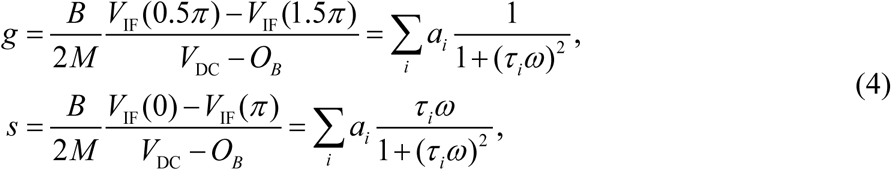

where *M/B* and ***O***_*B*_ are calibrated before measurements using the procedures detailed in Section S2.

### Image reconstruction and analysis

All image reconstructions and analyses were performed with our open-source *Instant-FLIM-Control* and *Instant-FLIM-Analysis* (Fig. S5) software developed for this work. Additional image processing and visualization were performed using ImageJ (US National Institutes of Health), Matlab (MathWorks), Imaris (Bitplane), or Illustrator (Adobe) software.

The raw intensity [Eq. (1)], *g*, and *s* [Eq. (4)] images from 2D, 3D, and 4D instant FLIM measurements were exported as 32-bit TIF files using the *Instant-FLIM-Control* program. Whereas the program allows the export of additional FLIM data, to reduce the consumption of the computer’s storage space, we exported only the raw intensity, *g*, and *s* images, which can be used to reconstruct other types of FLIM data. We used the “Correct 3D Drift” ImageJ plugin (*40*) to register the 4D raw images to correct for the sample drift during time-lapse measurements. To perform the image registration simultaneously on the three raw images, we created a color image by merging the raw intensity, *g*, and *s* images as its red, green, and blue channels, respectively; then, as the 3D image registration was performed on the red (intensity) channel, the other channels were registered simultaneously. The registered image was cropped to remove the empty voxels generated from the registration process, and its channels were split into three separate images, which correspond to the registered intensity, *g*, and *s* images.

The intensity, *g*, and *s* images were then imported to the *Instant-FLIM-Analysis* program to generate other types of FLIM data. Since the raw FLIM data usually have a low SNR (*11*), we provided a filtering option in the program, such that one could apply a median filter (3 × 3 or 5 × 5 kernels) or a smoothing filter on the raw images, one or multiple times, to increase the SNR; note that applying a median filter to the phasor images does not decrease the image resolution (*21*). In this work, we applied a 3 × 3 median filter three times on the raw phasor images for all *in vivo* measurements. The program could also uniformly scale the *g* and *s* images by a constant ratio to effectively zoom-in or zoom-out the phasor plot while keeping the calculated lifetime unaltered; a ratio between 0.7 to 1.0 was used in our measurements to post-compensate for the inaccuracy in system calibration and make sure that the majority of the phasor points fall inside the universal semicircle [(*g* − 0.5)^2^ + *s*^2^ = 0.25]. We then generated the phasor plot from the pre-processed *g* and *s* images. The phasor plot was a 2D histogram of the phasor components, *g* and *s*, where the magnitude (represented with a color map) of each grid (with a pre-defined size) in the 2D histogram represented the number of phasor points located inside that grid. Note that the phasor plot could be generated simultaneously with intensity and lifetime images in the *Instant-FLIM-Control* program. In the *Instant-FLIM-Analysis* program, however, the phasor plot could be further analyzed using phasor-based image segmentation techniques. To perform the phasor analysis, the program allowed the user to either (a) manually draw ROIs on the phasor plot and segment the pixels corresponding to the phasor points enclosed by the ROIs with different colors (*21*), or (b) automatically group the phasors into *K* clusters using the K-means clustering algorithm and segment the pixels corresponding to the phasor clusters with different colors (*22*); both techniques could segment the raw image into structures with different lifetime/phasor characteristics (Fig. S6). After the analysis described above, the raw lifetime image (gray-scale image with pixel value as lifetime), composite lifetime image (RGB image with pixel brightness as intensity, hue as lifetime), phasor plot (RGB image), and segmented images based on phasor labeling or phasor clustering techniques (RGB image) were generated and exported as 32-bit TIF files. For 3D images, the TIF files were imported into ImageJ and the maximized z-projections of the stacks were generated. For 4D measurements, the stacks were first converted to hyperstacks in ImageJ to facilitate the time-lapse analysis, and a time-lapse maximized z-projection sequence could be generated. We also used Imaris to visualize the 3D and 4D stacks (Fig. 2, A and C; Fig. S7 and Fig. S8; Movie S1-Movie S5, Movie S7) and enhance the qualitative information of FLIM by surface rendering the composite lifetime images (Fig. 5B; Movie S4).

To quantify the results of *in vivo* instant FLIM measurements (Fig. 3-Fig. 6), we extracted the raw lifetime as well as *g* and *s* values from the TIF files exported by the *Instant-FLIM-Analysis* program. Specifically, the files were opened in ImageJ, a single-plane ROI was drawn around the cell of interest, and the mean value was measured within that ROI. The plots representing the phasor locations of cells were generated using Illustrator (Fig. 3D, Fig. 5C). Using the *Instant-FLIM-Analysis* program, ROIs were drawn on the phasor plots to correspond with the pixels representing individual cells. Individual ROIs were drawn for each cell analyzed. The locations for each of these ROIs were then recapitulated within the universal phasor semicircle. These specific locations were determined by exporting each phasor plot with the drawn ROIs and determining the coordinates of the ROIs within the exported phasor plot image using ImageJ. The coordinates were then used to place representations of each cell in the proper location within the universal phasor semicircle in Illustrator. In Fig. 3-Fig. 6 separate statistical tests on *τ, g*, and *s* were performed to directly compare lifetime measurements between experimental groups. This method allows for determination of specific lifetime components which are different between samples or experimental conditions.

### Animal procedures and sample preparation

All animal studies were conducted in accordance with the University of Notre Dame Institutional Animal Care and Use Committee guidance. The acquisition parameters for all the images shown in this work are presented in Table S1.

Intravital imaging of the mouse brain was performed similarly to previously described (*41*). Briefly, Cx3cr1-GFP/+ mice (generated in-house by crossing Cx3cr1-GFP/GFP mice with wildtype C57Bl/6 mice) were anesthetized with ketamine and xylazine cocktail by intraperitoneal injection. The mouse’s head was secured and fixed in place using a stereotaxic instrument (Stoelting Co), and the skull was exposed with a midline scalp incision. A high-speed microdrill (Ideal Microdrill) equipped with a 0.7 mm burr (Fine Science Tools, 19007-09) was used to thin the skull to approximately 30 μm in thickness, using light sweeping motions to thin the skull gradually without applying significant pressure to the brain. In some instances, dental cement was used to border the thinned skull to form a bowl-like barrier to help contain water for imaging. Following surgery, the anesthetized mouse was administered a retro-orbital injection of 70 kDa Texas Red-Dextran (Invitrogen D1864) to enable fluorescent detection of blood vessels. The mouse was placed on the stage for imaging and its heart rate, respiration rate, and toe-pinch reflex were monitored periodically to ensure that it was fully anesthetized and not under duress.

The transgenic zebrafish line used in this study was *Tg(pu1:gfp)*. All embryos were generated from pairwise matings and raised at 28 °C until imaging. Stable, germline transgenic zebrafish were used for all experiments. Embryos of either sex were used for all experiments. At 4 dpf, the embryos were dechorionated and anesthetized with 3-amino-benzoic acid ester. After anesthetization, embryos were mounted on their ventral side in 0.8% low-melting-point agarose in 35 mm Petri dishes with glass bottoms. Following mounting, the embryos were imaged using the instant FLIM system described above.

To prepare a fixed mouse brain to image for select experiments, a living Cx3cr1-GFP/+ mouse was first deeply anesthetized with isoflurane. Next, the mouse’s chest cavity was opened to expose the heart, and the mouse was provided with a 100 μL injection of fixable Dextran 594 directly into the heart. After allowing the dextran to circulate for 1 minute, the mouse was perfused with 10 mL ice-cold 4% PFA to thoroughly fix the brain. Subsequently, the brain was extracted and fixed in 4% PFA overnight at 4 °C, and washed in 1x PBS, after which point it was ready for imaging.

MDA-MB-231-EGFP cells were cultured in DMEM High Glucose supplemented with 10% FBS and 1% Penicillin-Streptomycin in an incubator at 37 °C, 5% CO2. Prior to imaging, the cells were grown to about 80% confluence for the imaging experiments. During imaging, refrigerated or heated culture media were added to the cell culture to change the microenvironment temperature, which was simultaneously monitored by a thermometer placed in the media.

FluoCells prepared slide #1 (Invitrogen F36924) was used as a biological test slide to evaluate the performance of the instant FLIM system. This test slide contained bovine pulmonary artery endothelial (BPAE) cells labeled with multiple fluorescent dyes: the mitochondria were stained with MitoTracker Red CMXRos, the F-actin was labeled with Alexa Fluor 488 phalloidin, and the nuclei were labeled with DAPI.

Four lifetime standards, i.e., fluorophore solutions with known fluorescence lifetimes, were prepared as previously described (*26, 27*): 1 mM coumarin 6 in methanol (2.3 ns at 20 °C), 1 mM coumarin 6 in ethanol (2.4 ns at 20 °C), 1mM fluorescein in 0.1 M NaOH (4.0 ns at 20 °C), and 1 mM rhodamine B in water (1.7 ns at 20 °C). To prepare the standards, the coumarin 6 (Sigma-Aldrich 546283) was dissolved in methanol (VWR BDH2029) and ethanol (Millipore 818760), respectively; the fluorescein (Sigma-Aldrich F245-6) was dissolved in NaOH 1.0N in aqueous solution (VWR BDH7222); and the rhodamine B (Alfa Aesar A13572) was dissolved in water. The fluorophore mixtures were acquired by mixing the fluorescein and rhodamine B lifetime standards described above with different mole ratios.

## Supporting information

Movie S1. Instant FLIM imaging of a fixed mouse brain.

Movie S2. Through-skull in vivo instant FLIM imaging of intact mouse brains with adaptive optics.

Movie S3. Super-resolution instant FLIM imaging of a fixed mouse brain.

Movie S4. 4D in vivo instant FLIM imaging in an injured zebrafish brain.

Movie S5. 4D in vivo instant FLIM imaging in an injured mouse brain.

Movie S6. 4D in vivo lifetime and phasor imaging in injured mouse brains.

Movie S7. Phasor labeling and phasor clustering techniques applied to 4D in vivo instant FLIM stacks of an injured mouse brain.

## Supplementary Materials

Supplementary material for this article is available.

Section S1. Principle of instant FLIM

Section S2. System calibration

Section S3. SNR analysis of instant FLIM Section

S4. Instant FLIM with adaptive optics

Section S5. Super-resolution instant FLIM with GSOS

Fig. S1. Detailed diagram and photos of an instant FLIM system.

Fig. S2. Analog signal processing in instant FLIM.

Fig. S3. Instant FLIM system calibration using a lifetime standard.

Fig. S4. SNR analysis of instant FLIM and conventional FD-FLIM techniques.

Fig. S5. Overview of the *Instant-FLIM-Control* and *Instant-FLIM-Analysis* software.

Fig. S6. Instant FLIM with phasor labeling and phaser clustering techniques.

Fig. S7. Instant FLIM with adaptive optics enabling through-skull *in vivo* lifetime imaging of intact mouse brains.

Fig. S8. Super-resolution instant FLIM enabled by GSOS.

Table S1. Acquisition parameters for all data.

Table S2. Parts and price list for instant FLIM.

Movie S1. Instant FLIM imaging of a fixed mouse brain.

Movie S2. Through-skull *in vivo* instant FLIM imaging of intact mouse brains with adaptive optics.

Movie S3. Super-resolution instant FLIM imaging of a fixed mouse brain.

Movie S4. 4D *in vivo* instant FLIM imaging in an injured zebrafish brain.

Movie S5. 4D *in vivo* instant FLIM imaging in an injured mouse brain.

Movie S6. 4D *in vivo* lifetime and phasor imaging in injured mouse brains.

Movie S7. Phasor labeling and phasor clustering techniques applied to 4D *in vivo* instant FLIM stacks of an injured mouse brain.

## Author contributions

Y.Z. and S.S.H. conceived and coordinated the project, designed and built the microscope, wrote the control and image-processing software, and performed the imaging experiments. I.H.G. and S.Z. performed the mouse surgery, acquired the images, and analyzed the data. E.L.N. and C.J.S. prepared the zebrafish, performed the zebrafish imaging experiments, and analyzed the data. D.B. assisted with the microscope assembly and performed the temperature-dependent imaging experiment. S.S.H., S.Z., and C.J.S. supervised the study. All authors contributed to the writing and editing of the manuscript.

## Data and materials availability

The repositories of the open-source *Instant-FLIM-Control* (https://github.com/yzhang34/Instant-FLIM-Control.git) and *Instant-FLIM-Analysis* (https://github.com/yzhang34/Instant-FLIM-Analysis.git) software are hosted on GitHub and they are freely available for academic use. Detailed protocols describing how to use the two software packages are included within the repositories. All data needed to evaluate the conclusions in this paper are present in the paper and/or the Supplementary Materials. Additional data related to this paper may be requested from the authors.

## Supplementary Information

### Section S1. Principle of instant FLIM

We first assume the fluorescence sample has a single-exponential decay with an impulse response function *f* (*t*) = exp(−*t*/*τ*) /*τ*, where *τ* is the fluorescence lifetime and the function’s integral on the time domain (*t* ≥ 0) is normalized to one. A mode-locked Ti:sapphire laser which generates femtosecond pulses at 80 MHz is used as the excitation source. Since the femtosecond pulses (100 fs) are orders of magnitude shorter than the 80 MHz modulation period (12.5 ns), we can consider these pulses as a Dirac comb with a period of ***T*** = 12.5 ns. Mathematically, the excitation irradiance can be denoted as ***I*** (*t*) = ***P*** *δ*_***T***_ (*t*), where ***P*** is a coefficient related to the average power of the laser beam. Based on the quadratic nature of two-photon excitation, the fluorescence ***F*** (*t*) is proportional to the convolution of *I* ^2^(*t*) and *f* (*t*),

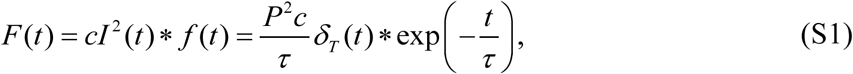

where *c* represents the fluorophore concentration. Since the fluorescence signal ***F*** (*t*) is periodic with an angular frequency of *ω* = 2*π*/***T***, it can be written as a Fourier series,

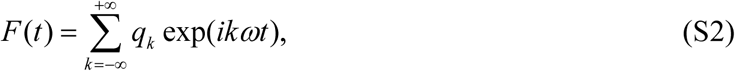

where the Fourier coefficients *q*_*k*_ are

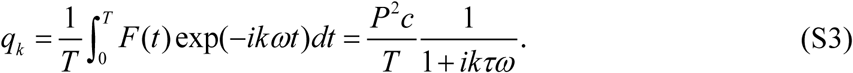

We separate the fluorescence signal ***F*** (*t*) detected by a PMT into DC and RF parts using a bias tee. The DC part is linearly related to *q*_0_ and can be written as

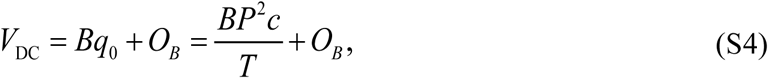

where ***B*** is the conversion loss from the bias tee’s RF&DC to DC ports, and ***O***_*B*_ is the offset that is invariant to the PMT signal variations. The RF part, on the other hand, is ∑_*k*≠0_ *q*_*k*_ exp(*ikωt*). Since 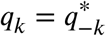, it can also be written as

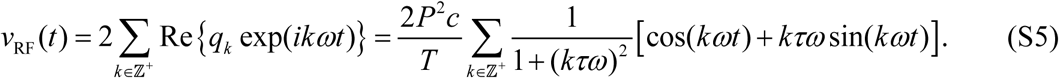

We use a homodyne detection method to extract the lifetime information from the first harmonic (1*ω*) of *v*_RF_(*t*). Specifically, we use RF mixers to mix the fluorescence signal with the 80 MHz reference signal from the Ti:sapphire laser. The reference signal is low-pass filtered and amplified such that it only contains the fundamental harmonic (1*ω*). For both the fluorescence and reference signals, a 4-way power splitter is used to split the signal to four paths. In instant FLIM, the four paths are operated independently and simultaneously during imaging acquisition. For each reference signal path, a phase shifter is utilized to introduce a voltage-controlled phase shift of *φ* (from 0 to 2*π*) to the signal. The reference signals are sent to the local oscillator (LO) ports of the mixers and can be denoted as

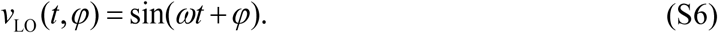

In instant FLIM, the four mixers take the PMT and reference signals to their RF and LO ports, respectively. In theory, the output signal on the intermediate frequency (IF) ports can be regarded as the product of the signals on its RF and LO ports; in practice, however, a DC offset due to circuit imbalance also exists in the output (*42*). We denote this DC offset as

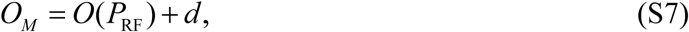

where ***O***(***P***_RF_) is a function of the RF signal’s power, ***P***_RF_, and *d* is a constant. The DC offset ***O***_***M***_ cannot be eliminated; it could be minimized by matching the powers of the RF and LO signals (*42*), but in our setup ***P***_RF_ is a variable related to the excitation power and the fluorophore concentration, so the powers of RF and LO do not match. Considering the existence of ***O***_***M***_, we model the output of the mixers as

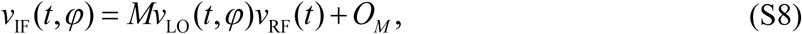

where *M* is the conversion loss from the mixer’s RF to IF ports. The mixers’ outputs are then low-pass filtered to eliminate all the harmonics; therefore, the filtered signals are pure DC voltages, as

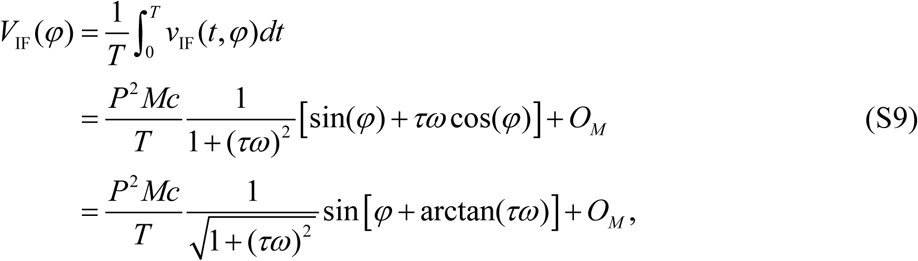

which are sampled by a DAQ card and used to calculate fluorescence lifetimes.

A calibration procedure described in Section S2 is required before measurement. When the calibration is complete, we can control the phase shifts *φ* by applying corresponding bias voltages to the phase shifters; we will also have the knowledge of ***M***/***B***, a ratio between the conversion losses of the mixer and the bias tee, that will be used for lifetime and phasor calculations. The instant FLIM measurement is performed by applying four different phase shifts, 0, 0.5*π, π* and 1.5*π*, respectively, to the four phase shifters on the reference signal paths. Based on Eq. (S9), the measured mixer outputs are

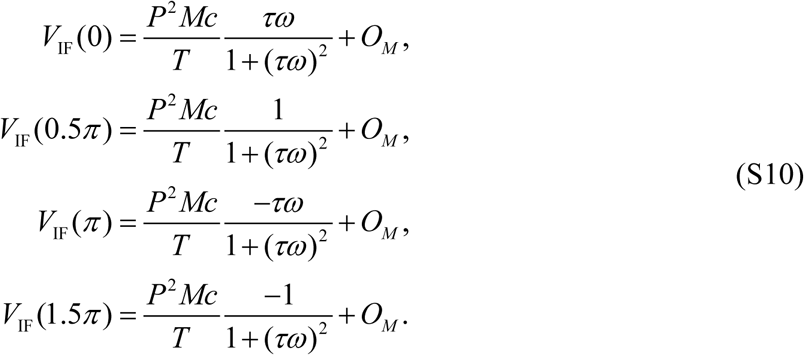

Then, by taking the differences among them, we have

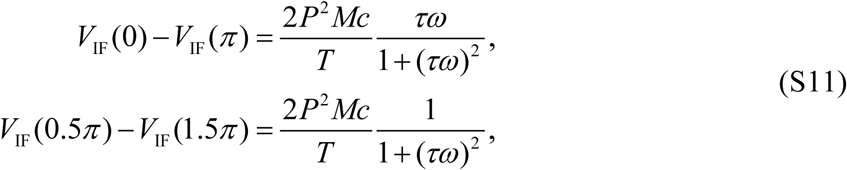

which are proportional to the imaginary and real parts of the first harmonic (1*ω*) Fourier coefficient of the fluorescence, *q*_1_, in Eq. (S3). Based on the measurements in Eq. (S11), it is easy to extract the fluorescence lifetime by 

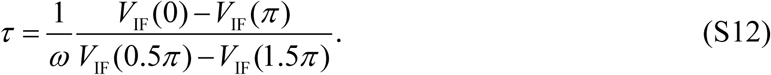

Next, we show that the instant FLIM method is also applicable for fluorophores with multi-exponential decays by employing the phasor plot approach (*21*). Considering that the fluorophore sample consists of multiple fluorophores with different lifetimes, *τ*_*i*_, the impulse response function becomes

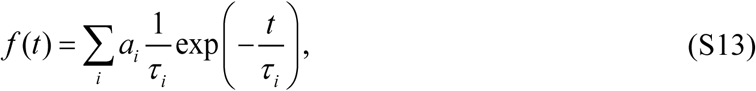

where *a*_*i*_ is the intensity weighted fractional contribution of the fluorophore with lifetime *τ*_*i*_, and ∑_*i*_ *a*_*i*_ = 1. Therefore, Eq. (S3) becomes

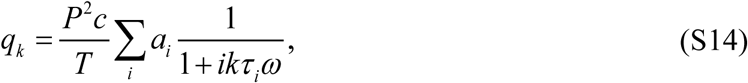

and Eq. (S11) changes to

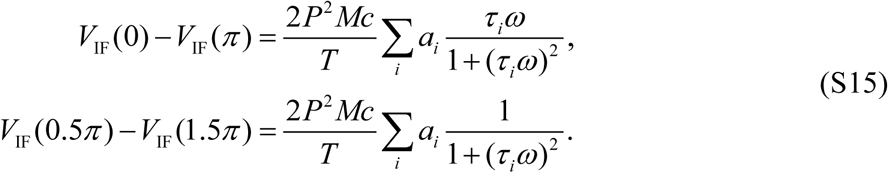

On the other hand, the DC signal from the bias tee, Eq. (S4), does not change when we assume a multi-exponential decay model. If we divide Eq. (S15) by 2(***V***_DC_ − ***O***_*B*_) in Eq. (S4) and the calibration coefficient ***M*** /***B***, we get

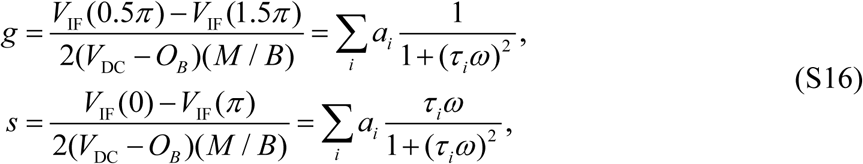

where *g* and *s* are the horizontal and vertical components (coordinates) of phasors on a phasor plot (*21*). From the phasor components, we can obtain the average fluorescence (phase) lifetime of the multi-exponential decay by

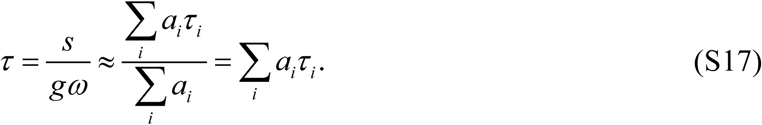

The average fluorescence lifetime alone cannot resolve the heterogeneity of multi-exponential decays, as different fluorophore compositions could result in the same average lifetime measurements. The phasor plot, on the other hand, can be used to resolve the heterogeneity because different fluorophore compositions can alter the phasor components even if the average lifetime might be unaltered.

### Section S2. System calibration

A fluorescence sample with a known lifetime (e.g., 10^−3^ M coumarin 6 in methanol with a lifetime of 2.30 ns (*27*)) is required for system calibration. The calibration should be repeated for each excitation wavelength that will be used. Once the calibration is complete, the system is ready for imaging, and no more calibration is required. First, the relations between the phase shifts *φ* and the phase shifters’ bias voltages ***V***_*b*_, i.e., *φ*(***V***_*b*_), should be calibrated for all four signal paths. Considering a fluorescence sample with a known lifetime 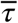 (e.g., 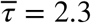 for coumarin 6 in methanol) and assuming that its concentration is 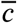, for each bias voltage ***V***_*b*_, Eq. (S9) becomes

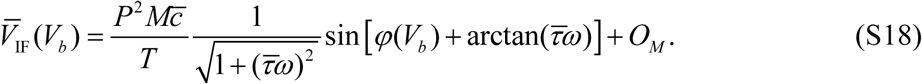

By changing ***V***_*b*_ and measuring 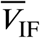, we can obtain a curve, 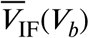, with maximal and minimal values:

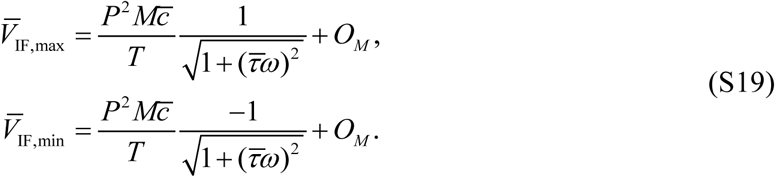

We then have

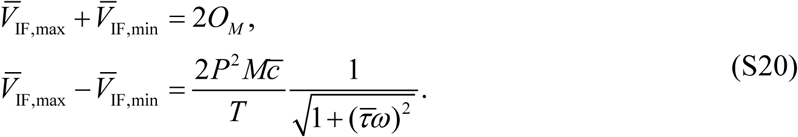

Thus, from Eq. (S18), we can extract the phase calibration curve, *φ*(***V***_*b*_), as

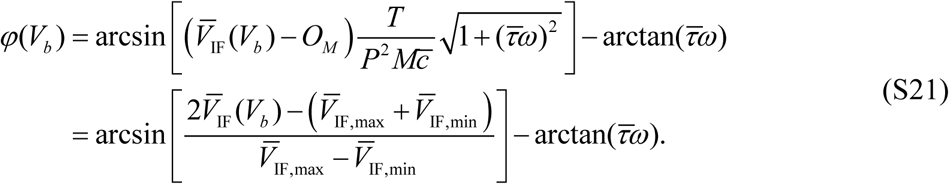

Therefore, an arbitrary phase shift *φ* between 0 to 2*π* can be introduced to the system by applying the corresponding voltage ***V***_*b*_ from the calibration curve *φ*(***V***_*b*_). Fig. S3 shows an example of 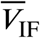 signals acquired from the four mixers’ IF ports, as well as the phase calibration curve *φ*(***V***_*b*_) calculated using Eq. (S21), during the system calibration using a coumarin 6 lifetime standard.

Second, the coefficient *M* /*B* used in Eq. (S16) and the offset ***O***_*B*_ in Eq. (S4) should be obtained during the calibration process. For the same fluorescence sample with a known lifetime 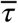, the DC signal from the bias tee, Eq. (S4), becomes

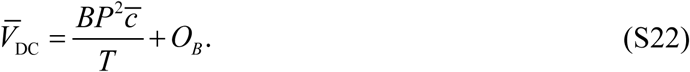

The offset ***O***_*B*_ can be obtained by turning off the laser beam (***P*** = 0) while measuring 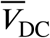. Therefore, based on Eqs. (S20) and (S22), we get

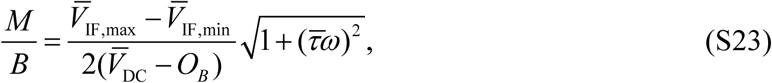

which is used for lifetime and phasor measurements.

Note that in Fig. S3 the four mixers have almost identical electrical properties as the calibration curves overlap with each other; therefore, it is valid to assume that the four mixers have identical conversion losses *M* and DC offsets ***O***_***M***_. However, while the conversion loss *M* is a design specification and can be guaranteed to be identical for the same type of mixers, there is no guarantee that the DC offsets ***O***_***M***_ are the same for all mixers (*42*). Thus, an additional calibration for the DC offset difference may be needed. We add a subscript *i* to denote the signals from the *i*-th mixer; the fluorescence signal from the *i*-th mixer’s IF port becomes

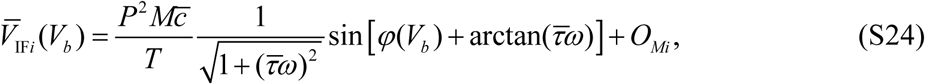

where the *i*-th DC offset is

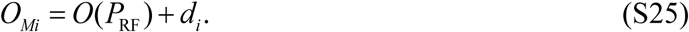

We found through experiments that ***O***(***P***_RF_), as a function of the RF signal’s power ***P***_RF_, was identical for all four mixers, and that the DC offset difference was due to *d*_*i*_. Therefore, we can compensate for such difference by calibrating the differences among *d*_*i*_. To do that, we turn off the laser beam (***P*** = 0) while measuring 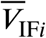 from the four mixers:

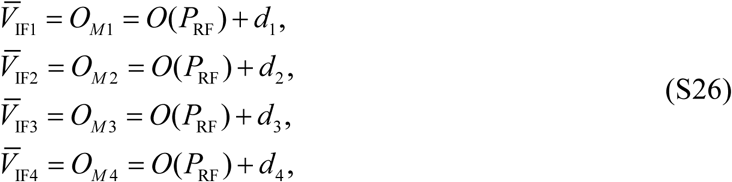

and we define two new calibration parameters, *d*_13_ and *d*_24_, as

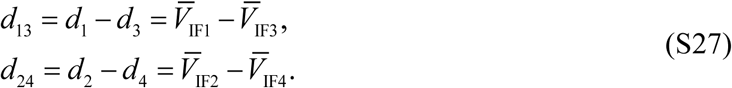

After the calibration, the DC offset differences among the mixers can be compensated by including *d*_13_ and *d*_24_ in FLIM measurements: Eq. (S12) becomes

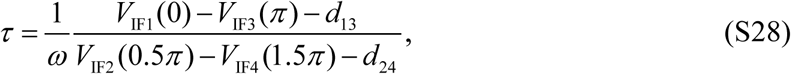

and finally, the phasor components [Eq. (S16)] are now

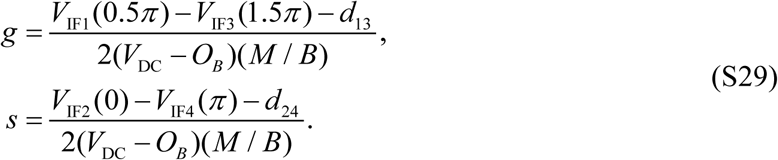

Note that all the calibration procedures described in this section can be performed easily and completed automatically with our open-source *Instant-FLIM-Control* program.

### Section S3. SNR analysis of instant FLIM

We analyze the SNR performance of instant FLIM and compare it with conventional FD-FLIM methods through Monte Carlo simulations and error-propagation analyses (*8, 11*). These FD-FLIM methods differ in the modulation waveforms of the excitation light, and consequently, the detected fluorescence signals are also different, as shown in Fig. S4A. These differences result in different sensitivity and SNR in lifetime measurements. Here, we use the *F*-value, i.e., photon economy, a widely used figure of merit in FLIM, to quantify and compare SNRs of FLIM measurements. It is defined as the ratio of the uncertainties in lifetime (*τ*) and intensity (*I*) measurements:

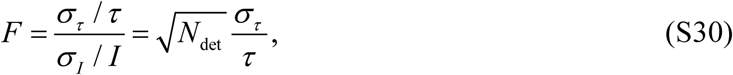

where we denote ***I*** = ***N***_det_, the number of photons detected in a measurement, and use the fact that ***N***_det_ is shot-noise-limited, i.e., Poisson distributed, such that 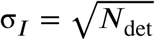. Note that the shot-noise-limited performance is achieved using high-sensitivity PMTs and low-noise amplifiers in a FLIM system. Since the *F*-value quantifies the uncertainty in lifetime measurements, a smaller *F* means that the lifetime is measured more accurately, hence a better SNR performance. Whereas a smaller *F* is desired, it is limited to ***F*** > 1 due to shot noise; ***F*** = 1 only exists in an ideal shot-noise-limited FLIM system.

We use Monte Carlo simulations and error-propagation analyses to acquire the *F*-values of instant FLIM and conventional FD-FLIM methods. The Monte Carlo simulations are performed by dividing each modulation period *T* (12.5 ns for 80 MHz modulation) into *L* time units Δ*t* (*8, 11*). In each Δ*t*, a random number, *r*, that is uniformly distributed in [0, 1] is generated and compared with the probability density described by the product of the fluorescence signal ***F***(*t*) and the time unit Δ*t*. If *r* is larger than ***F***(*t*)Δ*t*, the simulation considers that a fluorescence photon is emitted and recorded by the detector. In the analysis of instant FLIM, ***F***(*t*) is identical to Eq. (S1), while in other cases, ***F***(*t*) are analytically calculated by convolving the corresponding excitation signal (Fig. S4A) with the impulse response function, *f* (*t*). A modulation period *T* is completed after the simulation goes through all Δ*t* in that period; then, another *T* is simulated likewise, and the emitted photons are cumulatively recorded by the simulation. For a single lifetime measurement, ***N*** modulation periods are simulated, and the lifetime *τ* is calculated based on the FD-FLIM technique in use. This lifetime measurement process is then repeated for 1,000 times to obtain the statistical properties (mean and variance) and consequently, the *F*-value, of the lifetime measurement. A diagram summarizing the Monte Carlo simulation process is shown in Fig. S4B.

The analytical error-propagation analysis is applicable when the lifetime can be written in the following form:

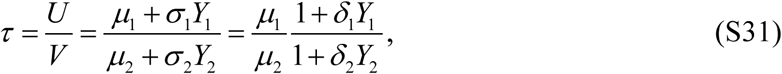

where *U* and *V* are the random variables used in lifetime calculation with means, *μ*_1_, *μ*_2_, standard deviations, *σ*_1_, *σ*_2_, and coefficients of variation, *δ*_1_ = *σ*_1_/*μ*_1_, *δ*_2_ = *σ*_2_/*μ*_2_; ***Y***_1_ and ***Y***_2_ are auxiliary random variables with zero means and unity variances. In practice, 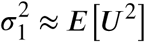 and 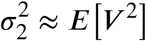 as ***E***[***U***]^2^ and ***E***[***V***]^2^ are negligible compared to ***E***[***U***]^2^ and ***E***[***V***]^2^. If the moments of third and higher orders are omitted, Eq. (S31) can be expanded as

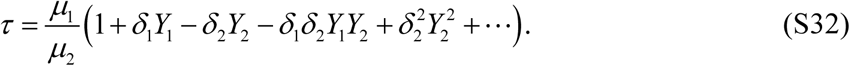

We then get the expected value of *τ* as

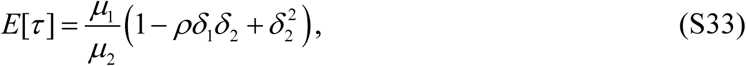

where *ρ* = ***E***[***Y***_1_***Y***_2_] is the correlation coefficient of *U* and *V*. We also have

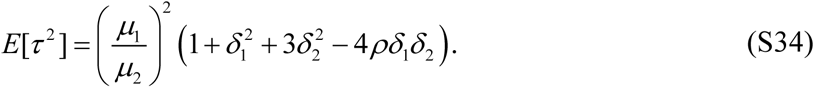

Therefore, the variance of *τ* can be calculated as

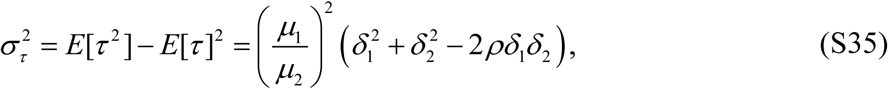

and the *F*-value can be calculated with Eq. (S30).

Based on the principle of instant FLIM in Section S1, the fluorescence signal ***F***(*t*) can be analytically calculated as

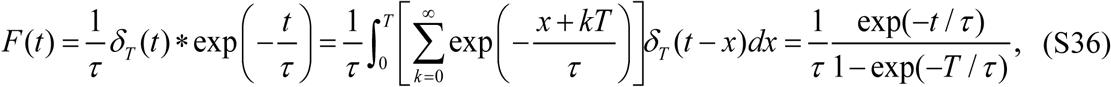

where we scale the excitation power and the fluorophore concentration down to ***P*** = 1 and *c* = 1 such that on average only one photon is emitted during a modulation period *T*, i.e., 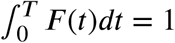. Thus, ***F*** (*t*) can be regarded as the probability density function of detecting a photon, and the mathematical expectation of a random variable *X* related to detecting a photon is

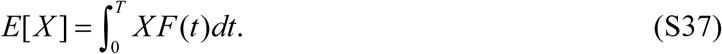

We denote the random processes corresponding to the four mixers’ outputs [Eq. (S9)] in instant FLIM as ***X***_1_, ***X***_2_, ***X***_3_ and ***X***_4_. With Eqs. (S6) and (S8), Eq. (S9) becomes

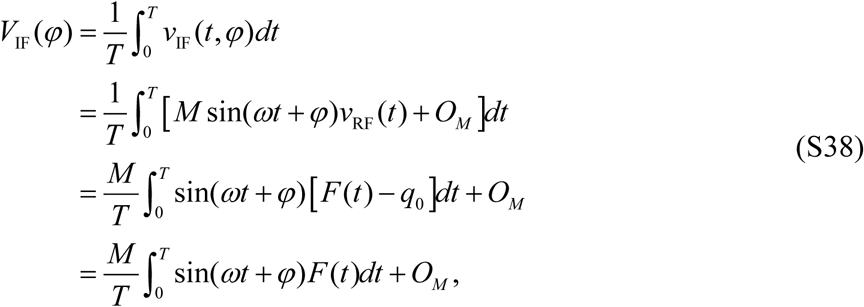

where we use the fact that *v*_RF_(*t*) and *q*_0_ are the RF and DC parts of ***F*** (*t*) and therefore *v*_RF_(*t*) can be written as *v*_RF_(*t*) = ***F*** (*t*) − *q*_0_. With Eqs. (S37) and (S38), we can write the random processes as

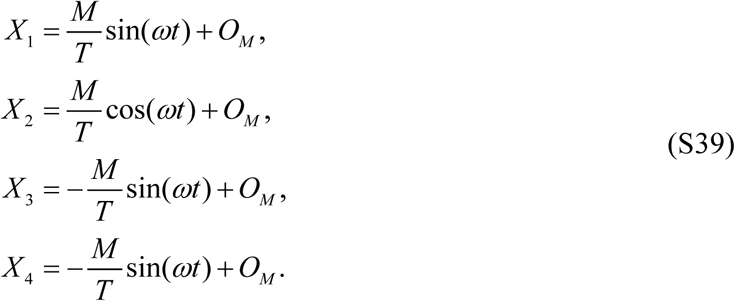

Since Eq. (S12) follows the form of Eq. (S31), we have ***U*** = ***V***_IF_(0) − ***V***_IF_(π) and ***V*** = *ω****V***_IF_(0.5π) − ***V***_IF_(1.5π). Considering that *N*_det_ photons are detected in a lifetime measurement, based on Eqs. (S3) and (S39), we get the means of *U* and ***V*** as

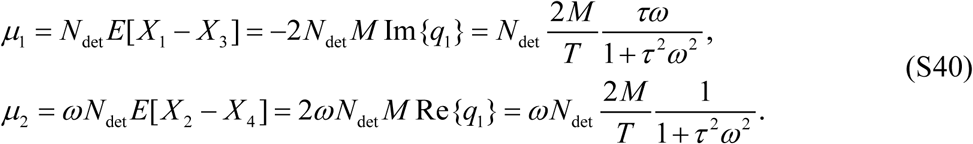

The variances and correlation coefficients of *U* and *V* can also be obtained as

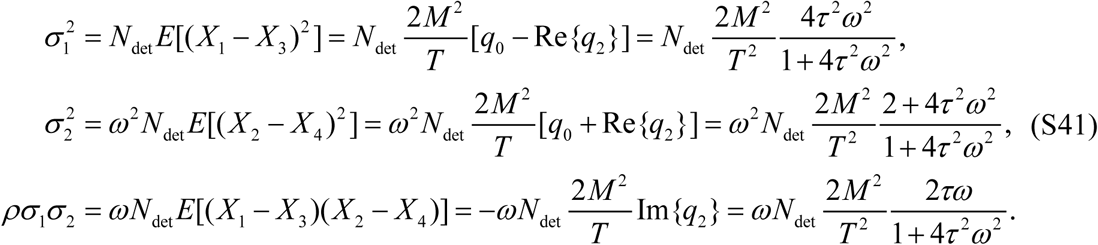

Then, from Eq. (S35), the standard deviation of *ဴ* is

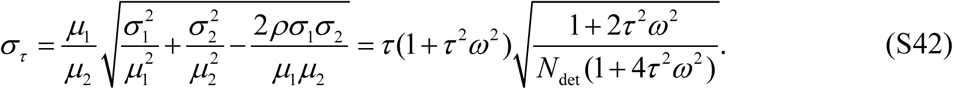

Consequently, we get the *F*-value of instant FLIM as

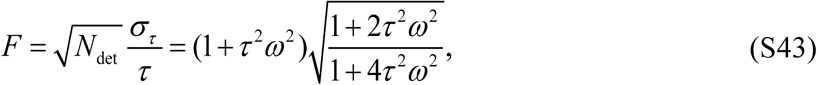

which is identical to the *F*-value of a two-photon (2P) FD-FLIM when the excitation is modulated as a Dirac comb (*11*); this is expected as instant FLIM uses a femtosecond pulse excitation in a 12.5 ns (80 MHz) modulation period, which can be seen as a Dirac comb.

We compare the SNR performances of instant FLIM and conventional FD-FLIM methods (*11*), including 2P FD-FLIM with sinusoidal (1.0, 0.5 modulation degrees) and periodic square wave (0.2, 0.5 duty cycles) modulations, as well as one-photon (1P) FD-FLIM with sinusoidal modulation (1.0 modulation degree), by plotting their simulated (symbols) and analytical (curves) *F*-values in Fig. S4C. Since the *F*-value is only dependent on the product of the modulation frequency and the fluorescence lifetime (*8*), i.e., *ωτ*, and the modulation frequency for instant FLIM is fixed at 80 MHz, for a fair comparison, we fix the modulation frequency of all the methods to 80 MHz and vary the lifetime from 0 to 4.5 ns when we investigate the *F*-values. The analytical results are in good agreement with the ones obtained from Monte Carlo simulations. Our instant FLIM method has the smallest *F*-value (***F*** = 1), which is the shot noise limit and the best possible SNR performance a FLIM method can achieve. In comparison, the 2P sinusoidal (1.0, 0.5 modulation degrees) and periodic square wave (0.2, 0.5 duty cycles) modulations have their best SNR performances at ***F*** = 2.62, ***F*** = 4.07, ***F*** = 1.44, and ***F*** = 2.89, respectively; the conventional 1P sinusoidal (1.0 modulation degree) case has its best *F*-value at ***F*** = 3.67. To illustrate the SNR difference among these FD-FLIM methods, we select four targeted lifetime values (0.375 ns, 1.625 ns, 2.875 ns, and 4.125 ns) and plot the traces of 1,000 measurements of their values using different FD-FLIM methods from the Monte Carlo simulations (Fig. S4D). A lifetime histogram and the standard deviation (*σ*_*τ*_) of the 1,000 lifetime measurements are plotted next to each trace. In all four cases, the instant FLIM measurements have the smallest *σ*_*τ*_ compared to conventional FD-FLIM methods. In summary, through Monte Carlo simulations and error-propagating analyses, instant FLIM has been demonstrated to be superior in SNR performance compared with conventional FD-FLIM techniques; in some cases, instant FLIM can approach the shot noise limit (***F*** = 1), the best possible SNR performance of a FLIM measurement. This is achieved by the efficient utilization of the 80 MHz femtosecond pulsed laser in instant FLIM.

## Section S4. Instant FLIM with adaptive optics

Adaptive optics (AO) can be employed in instant FLIM to reduce the optical aberrations induced by the imaging system or the sample, and potentially improve the imaging quality and extend the penetration depth (*23*). We implemented AO as an optional module in instant FLIM by introducing a deformable mirror to the setup. To reduce the system complexity and cost, we employed sensorless AO, in which a wavefront sensor was not required, and indirect wavefront sensing based on an optimization metric was used. Using the optimization algorithms described below, the AO module iteratively adjusts the parameters of the deformable mirror to improve the image-based optimization metric.

Depending on the user’s choice in the *Instant-FLIM-Control* program, the optimization metric, *V*, can be the mean pixel value or the normalized variance of the image, where the normalized variance is defined as the variance of all the pixel values divided by the square of the mean pixel value. We employed Zernike modes to describe the optical aberrations as well as the aberration compensation applied to the deformable mirror. The optimization algorithms are performed in an *N*-dimensional aberration space generated by a base:

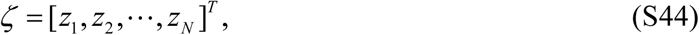

where *z*_1_, *z*_2_, …, *z*_*N*_ are Zernike polynomials. The deformable mirror (Thorlabs DMP40-P01) we used supports up to 12 Zernike modes 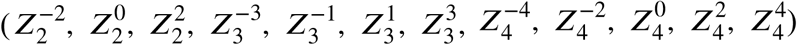 in the optimization. The aberration compensation, Φ, can then be described in the aberration space spanned by *ζ*:

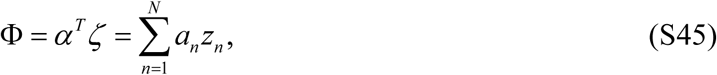

where

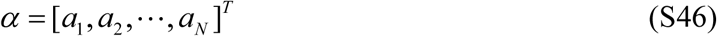

are the Zernike coefficients in which *a*_*j*_ represents the contribution of *z*_*j*_ to Φ. In practice, an optimization algorithm iteratively monitors the image-based metric *V* and updates the Zernike coefficients *α* that control the wavefront shaping elements (mirror segments) on the deformable mirror; after multiple iterations, if the AO optimization is successful, the metric *V* will be increased and the imaging quality will be improved.

In the *Instant-FLIM-Control* program, we use three AO algorithms to optimize *V* and imaging quality. The first one is a simple max search algorithm that sequentially searches through each Zernike coefficient, from *a*_1_ to *a*_*N*_, until a maximized *V* is found for each one of them. For example, to find the optimal value for *a*_*j*_, the algorithm keeps all the other Zernike coefficients unaltered while changing *a*_*j*_ from −1 to 1 with a step size defined by the user, e.g., 0.2. This is valid due to the orthogonality of the Zernike modes. The optimal *a*_*j*_ is found at the value where *V* is maximized from −1 < *a*_*j*_ < 1. This algorithm is easy and straightforward, but it is time-consuming: for a step size of 0.2, full optimization of 12 Zernike modes requires the acquisition of 120 raw images. However, this is also the most robust and reliable algorithm, as it guarantees that a global optimum or its proximity can be found, whereas the other algorithms are likely to find a local optimum.

The second algorithm is a quadratic search algorithm (*28*) which approximates the metric *V* of the aberration Φ as a quadratic function of the Zernike coefficients *α*:

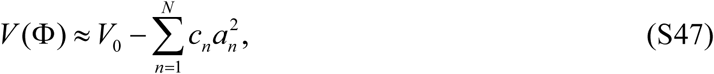

where *V*_0_ and *c*_*n*_ are constants and the metric has a paraboloidal shape around its maximum. The goal of the optimization algorithm is to find the coordinates of α that maximizes *V*. This process can be decomposed into *N* independent one-dimensional parabolical optimization problem for each Zernike coefficient *a*_*n*_, as Eq. (S47) can be written as

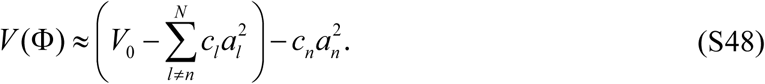

The optimal *a*_*n*_ that maximizes *V* can then be calculated by three measurements of *V*, i.e., ***V*** (Φ_0_), ***V*** (Φ_0_ + *bz*_*n*_), and ***V*** (Φ_0_ − *bz*_*n*_) corresponding to *a*_*n*_ = *a*_*n*0_, *a*_*n*_ = *a*_*n*0_ + *b*, and *a*_*n*_ = *a*_*n*0_ − *b*, respectively, where Φ_0_ is the initial aberration, *a*_*n*0_ is the initial value of *a*_*n*_ (*a*_*n*0_ = 0 by default), and *b* is the bias amplitude used to search the parabolic maximum. With the three metrics obtained from the three raw images, the optimal *a*_*n*_, i.e., *a*_*nn*_, can be estimated using parabolic maximization as

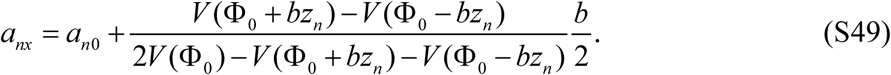

In the algorithm, this one-dimensional optimization process is repeated for *N* times to calculate the optimal Zernike coefficients *a*_1*n*_, *a*_2*n*_, …, *a*_*Nn*_ that jointly maximizes *V*. Whereas each process requires three measurements, the metric for the initial image ***V*** (Φ_0_) is common; therefore, to correct for *N* Zernike coefficients, the algorithm merely requires 2*N* + 1 raw images. For our deformable mirror supporting 12 Zernike modes, the optimization needs only 25 raw images. Compared to the other AO optimization algorithms, the quadratic search one is the fastest and requires the least number of raw images to be captured. However, it is not always reliable as the approximation assumption in Eq. (S47) is only accurate for small aberration amplitudes, which may not be satisfied in practice. Therefore, this algorithm is recommended only if the speed is an essential requirement for AO optimization.

The third algorithm in the program is based on the stochastic parallel gradient descent (SPGD) method (*29*). In each SPGD iteration, after the metric ***V*** is acquired from a raw image, the Zernike coefficients in *α* are perturbed in parallel by a small non-zero stochastic value. In our program, the value is randomly chosen from [−0.05, −0.025, 0.025, 0.05]. During the *m*-th iteration, the Zernike coefficients applied to the *m+1*-th iteration is calculated as

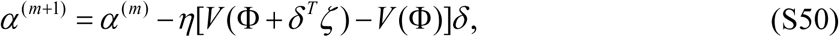

where the superscripts in brackets are the indices of the iteration, *η* is the learning rate, Φ is the aberration before the iteration, *ζ* is the Zernike base described in Eq. (S44), and *δ* is an array consisting of the non-z ero stochastic values. The learning rate *η* is negative in our case (−0.01 by default) as the optimization is to maximize *V*. If *η* is chosen properly, the metric *V* will be increased after each iteration. The algorithm is stopped when a maximal *V* is found or the specified steps of iterations are reached. Compared to the previous algorithms, the SPGD method updates the Zernike coefficients in parallel and it could acquire the optimal coefficients more accurately. However, due to the stochastic nature of the method, the algorithm usually takes much more iterations, hence imaging time, than the other methods, and it could also easily fall into a local maximum. As a complement to the other algorithms, the SPGD method is recommended when high precision is preferred in determining theptoimal Zernike coefficients.

To demonstrate the performance of instant FLIM when the AO module is added, we acquired through-skull 3D FLIM stacks of the intact brain in a living mouse with and without AO. We used the simple max search algorithm for the AO optimization. With AO, the instant FLIM system was able to generate 3D intensity, lifetime, and phasor labeled stacks of the living mouse brain, through the skull, with a penetration depth up to 300 μm (Fig. S7 and Movie S2). Note that the excitation power of the laser was gradually increased from 8.53 mW to 21.13 mW during the imaging; the penetration depth limit was achieved when further increasing the power could not increase the signal level. We then imaged the animal with the same imaging condition but without the AO module. Specifically, we added a folding mirror in front of the AO module to reflect the laser beam directly into the two-photon microscope. As shown in Fig. S7, when the AO module was not used, we could only achieve a penetration depth of 130 μm. In comparison, the penetration depth for instant FLIM with a well-optimized AO module (300 μm) was more than double of that without AO.

### Section S5. Super-resolution instant FLIM with GSOS

We show that the instant FLIM system can be used to generate super-resolution FLIM images based on the recently demonstrated generalized stepwise optical saturation (GSOS) technique (*24*). By linear combining *M* FLIM images obtained with different excitation powers in the complex domain, a new FLIM image with a 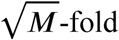 increase in spatial resolution can be obtained. Because the linear combination in GSOS is applied to the complex Fourier coefficients, *q*_*k*_, of the fluorescence, *F* (*t*), we rewrite Eq. (S5) such that *q*_*k*_ is explicitly shown in the equation:

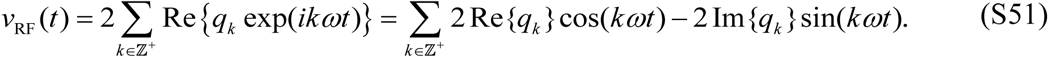

Following the analog signal processing in instant FLIM, the output signals of the mixers [Eq. (S9)] become

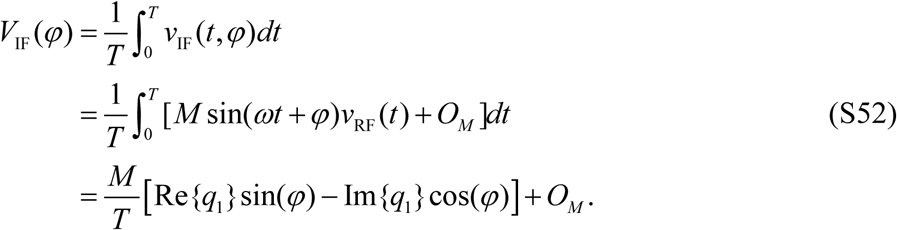

Consequently, after introducing the phase shifts, the measured mixer outputs [Eq. (S10)] are

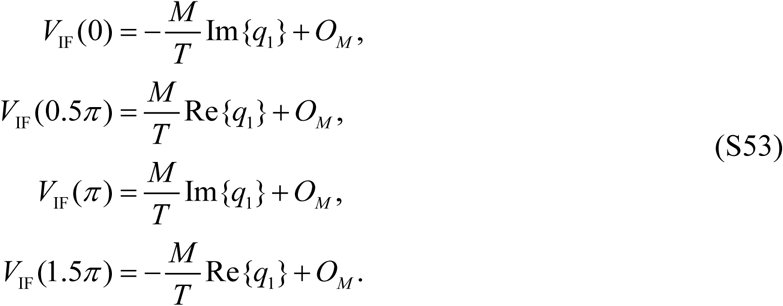

Then, the differences among them [Eq. (S11)] are directly proportional to the real and imaginary parts of the Fourier coefficient:

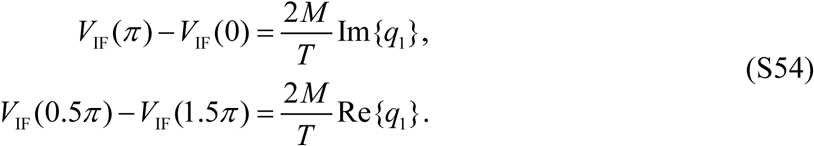

Therefore, the magnitude and phase of the complex Fourier coefficient, *q*_1_, can be extracted as

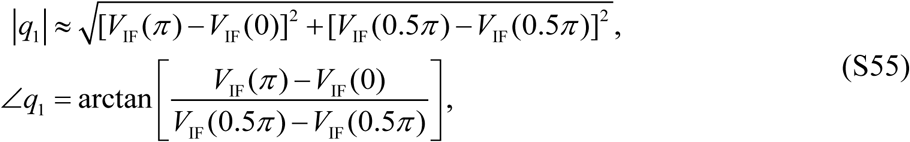

which are then combined as a complex value, *q*_1_ = |*q*_1_| exp(*i*∠*q*_1_).

In GSOS with *M* steps (M-GSOS), the measurement is repeated for *M* times with *M* different excitation powers, ***I***_01_, ***I***_02_, …, ***I***_0***M***_, where ***I***_01_ < ***I***_02_ < … < ***I***_0***M***_. Hence *M* images consisting of these complex Fourier coefficients, *q*_1,1_, *q*_1,2_, …, *q*_1,***M***_, are obtained based on the measurements described above [Eqs. (S53)-(S55)]. The GSOS linear combination is then applied to these complex images,

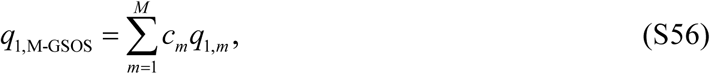

where the linear combination coefficients, *c*_*m*_, are calculated based on the excitation powers, *I*_01_, *I*_02_, …, *I*_0***M***_. Here, we give the general expression to calculate *c*_*m*_:

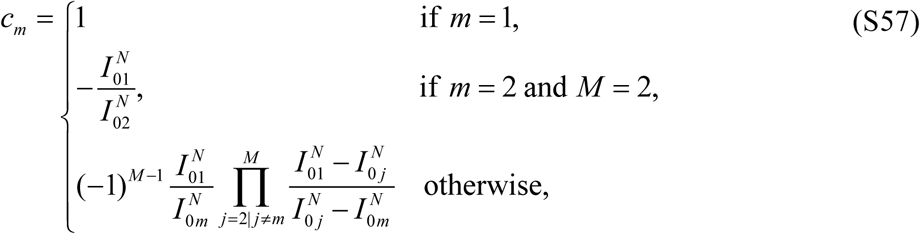

where *N* is the number of excitation photons needed for a fluorophore to emit one photon (***N*** = 1 for 1PEF, ***N*** = 2 for 2PEF). The result of the GSOS linear combination is a new image (*q*_1,M-GSOS_) with complex pixel values, where its magnitude, *q*_1,M-GSOS_, is a super-resolution image with a 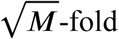 increase in resolution over the diffraction limit, and its phase, ∠*q*_1,M-GSOS_, is identical to that of the original Fourier coefficient, ∠*q*_1_ (*24*). From the phase, a fluorescence lifetime image can be obtained by

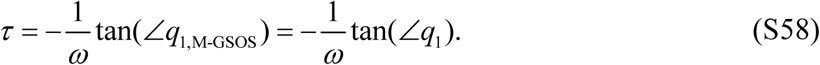

As a result, by linear combining *M* complex images obtained with an instant FLIM system using the GSOS principle, we can obtain a super-resolution FLIM image where its intensity, |*q*_1,M-GSOS_|, is enhanced by a factor of 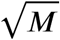 and its lifetime information, *τ* = − tan∠*q*_1,M-GSOS_ /*ω*, is preserved.

Instant FLIM can be used to generate super-resolution 3D FLIM stacks. Fig. S8 shows a side-by-side comparison between a diffraction-limited (DL) and a super-resolution (two-step GSOS) 3D FLIM stacks of a fixed mouse brain. The DL stack was acquired with an instant FLIM system at an excitation power of 12.31 mW. The two-step GSOS stack, on the other hand, was generated by linear combining two DL instant FLIM stacks, *q*_1,1_ and *q*_1,2_, measured at ***I***_1_ = 12.31 mW and ***I***_2_ = 13.44 mW, respectively. The GSOS linear combination coefficients were calculated according to Eq. (S57) (*N* = 2 for 2PEF): *c*_1_ = 1 and 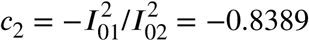. The two-step GSOS stack, *q*_1,2-GSOS_, was then generated based on Eq. (S56), i.e., *q*_1,2-GSOS_ = *c*_1_*q*_1,1_ + *c*_2_*q*_1,2_ = *q*_1,1_ − 0.8389*q*_1,2_; its intensity, *q*_1,2-GSOS_, and fluorescence lifetime, *τ* = − tan∠*q*_1,2-GSOS_ /*ω*, are plotted as brightness and hue, respectively, to construct the 3D FLIM stack. The resolution improvement of the GSOS stack compared to the DL stack can be seen in Fig. S8 (Movie S3), where the line profiles of the normalized intensity values at the positions of the white arrowheads show the features that are better resolved in the GSOS stack.

**Fig. S1.**
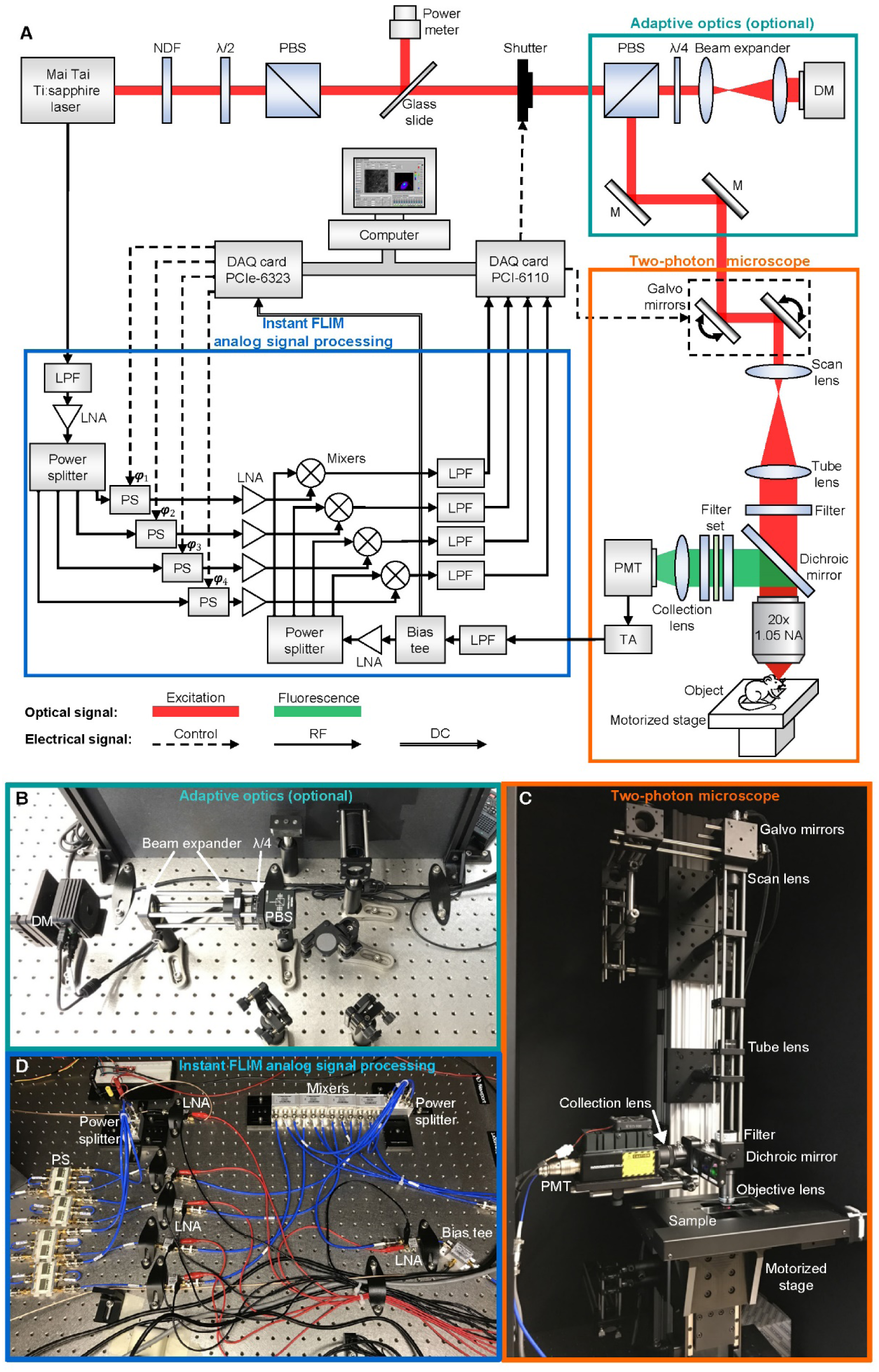
Detailed diagram and photos of an instant FLIM system. (**A**) Detailed diagram of the system configuration. NDF, neutral density filter; λ/2, half-wave plate; PBS, polarizing beam splitter; λ/4, quarter-wave plate; DM, deformable mirror; M, mirror; PMT, photomultiplier tube; TA, transimpedance amplifier; DAQ, data acquisition; LPF, low pass filter; LNA, low-noise amplifier; PS, phase shifter. Three sets of components are grouped together and highlighted as system modules. The two-photon microscope module is a conventional two-photon laser scanning microscope setup; the optional adaptive optics module can improve the imaging quality and penetration depth; the analog signal processing module is the essential part of an instant FLIM system. (**B**) Photo of an implementation of the optional adaptive optics module. (**C**) Photo of an implementation of the conventional two-photon microscope module. (**D**) Photo of an implementation of the instant FLIM analog signal processing module.

**Fig. S2.**
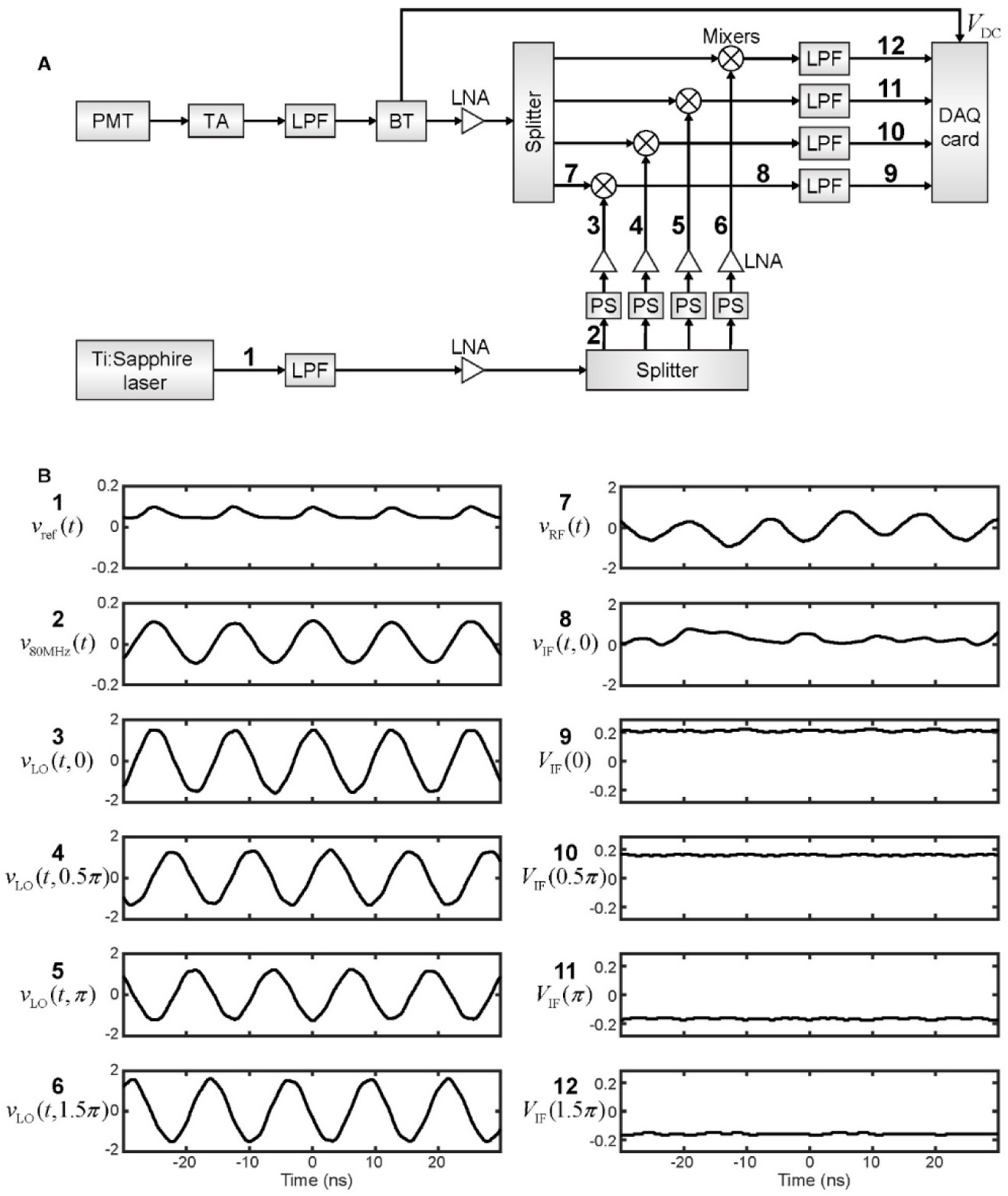
Analog signal processing in instant FLIM. (**A**) Diagram showing how analog signals are processed in instant FLIM. The signals are labeled with numbers and their typical waveforms are shown in (B). (**B**) Waveforms of the analog signals under real experimental conditions: excitation wavelength, 920 nm; sample, coumarin 6 in methanol; excitation power, 3.6 mW. These signals are: 1, *v*_ref_ (*t*), the reference signal from the mode-locked Ti:sapphire laser; 2, *v*_80MHz_(*t*), the pure 80 MHz signals after low pass filtering, amplifying, and splitting the reference signal; 3-6, *v*_LO_(*t*, φ), the phase-shifted reference signals directed to the mixers’ LO ports; 7, *v*_RF_(*t*), the low pass filtered, amplified, and split PMT signals directed to the mixers’ RF ports; 8, *v*_IF_(*t*, φ), one of the mixed signals generated on the mixers’ IF ports; 9-12, ***V***_IF_(φ), the low pass filtered (DC) mixed signals that are measured by a DAQ card. Unit, volts.

**Fig. S3.**
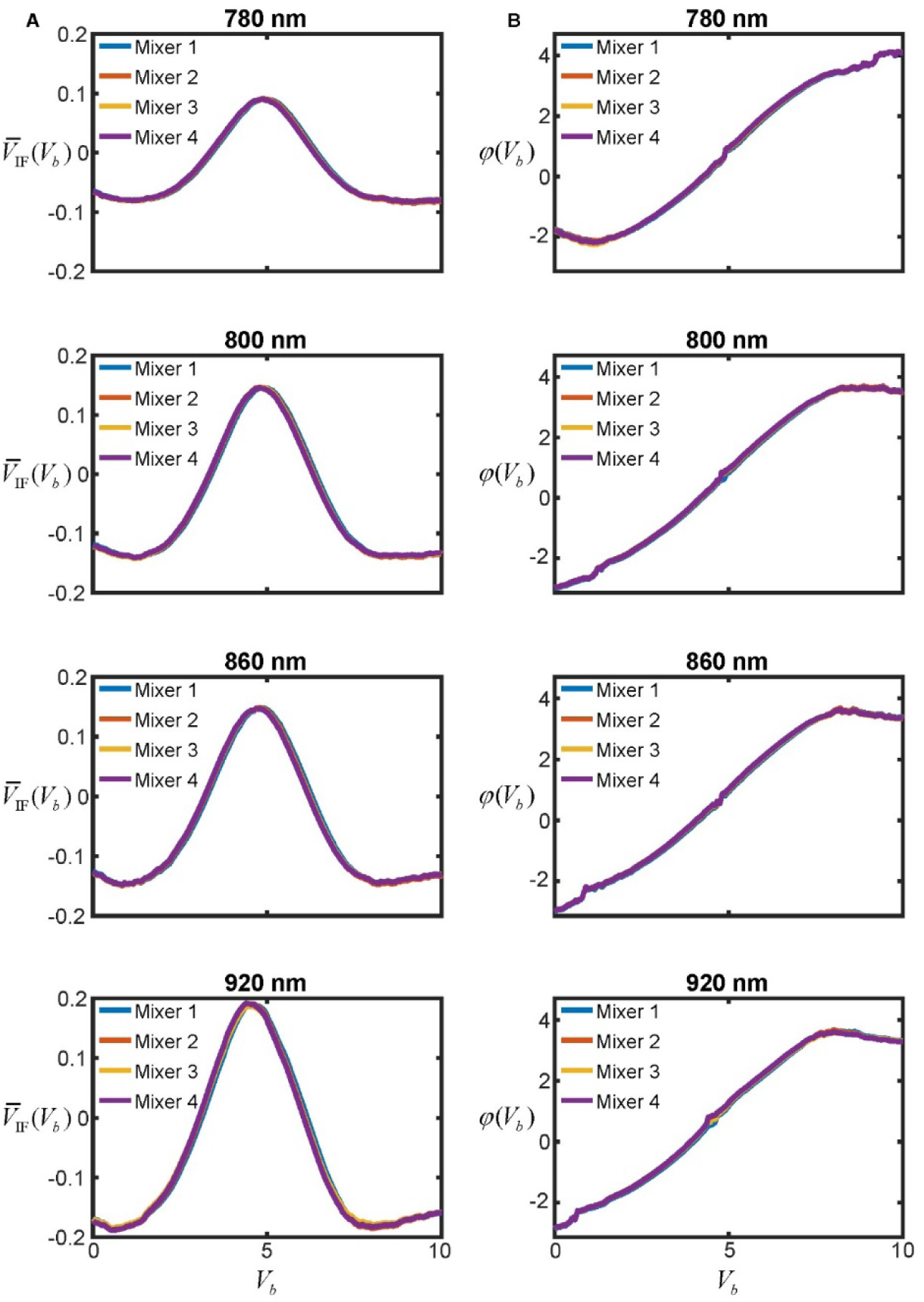
Instant FLIM system calibration using a lifetime standard. (**A**) Raw calibration curves acquired from a lifetime standard (1 mM coumarin 6 in methanol) by changing the bias voltage (***V***_*b*_, stepping from 0 to 10V) applied to the four phase shifters and measuring the voltages 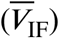 on the four mixers’ IF ports. A set of calibration curves should be measured for each excitation wavelength that will be used for imaging. Here we show the curves for excitation wavelengths at 780 nm, 800 nm, 860 nm, and 920 nm. Unit, volts. (**B**) Phase calibration curves, *φ*(***V***_*b*_), generated using Eq. (S21) based on the raw data in (A). Unit, radians.

**Fig. S4.**
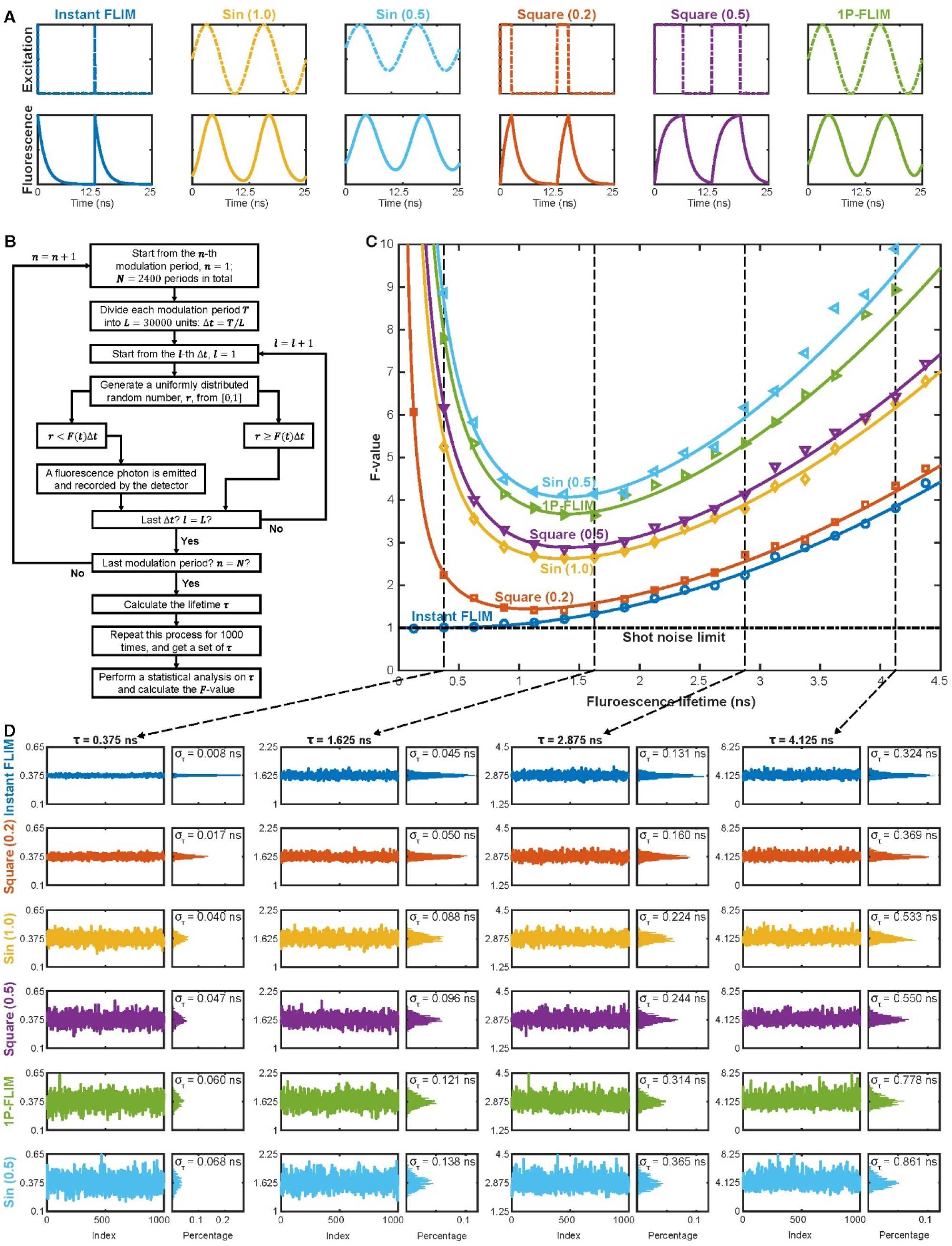
SNR analysis of instant FLIM and conventional FD-FLIM techniques. (**A**) Excitation and fluorescence waveforms of the six FD-FLIM techniques in comparison: instant FLIM, this work; Sin (1.0), two-photon (2P) sinusoidal (1.0 modulation degree) modulation; Sin (0.5), 2P sinusoidal (0.5 modulation degree) modulation; Square (0.2), 2P periodic square wave (0.2 duty cycle) modulation; Square (0.5), 2P periodic square wave (0.5 duty cycle) modulation; 1P-FLIM, one-photon (1P) sinusoidal (1.0 modulation degree) modulation. Modulation period, 12.5 ns (80 MHz). (**B**) Diagram summarizing the Monte Carlo simulation process. (**C**) *F*-values as a function of fluorescence lifetimes with modulation frequency fixed at 80 MHz for different FD-FLIM methods. The symbols and curves are results of the numerical Monte Carlo simulation and the analytical error-propagation analysis, respectively. (**D**) Simulated lifetime measurements extracted from four different lifetime values in the Monte Carlo simulations for different FD-FLIM methods. The histograms as well as the standard deviations of the lifetime measurements are also shown.

**Fig. S5.**
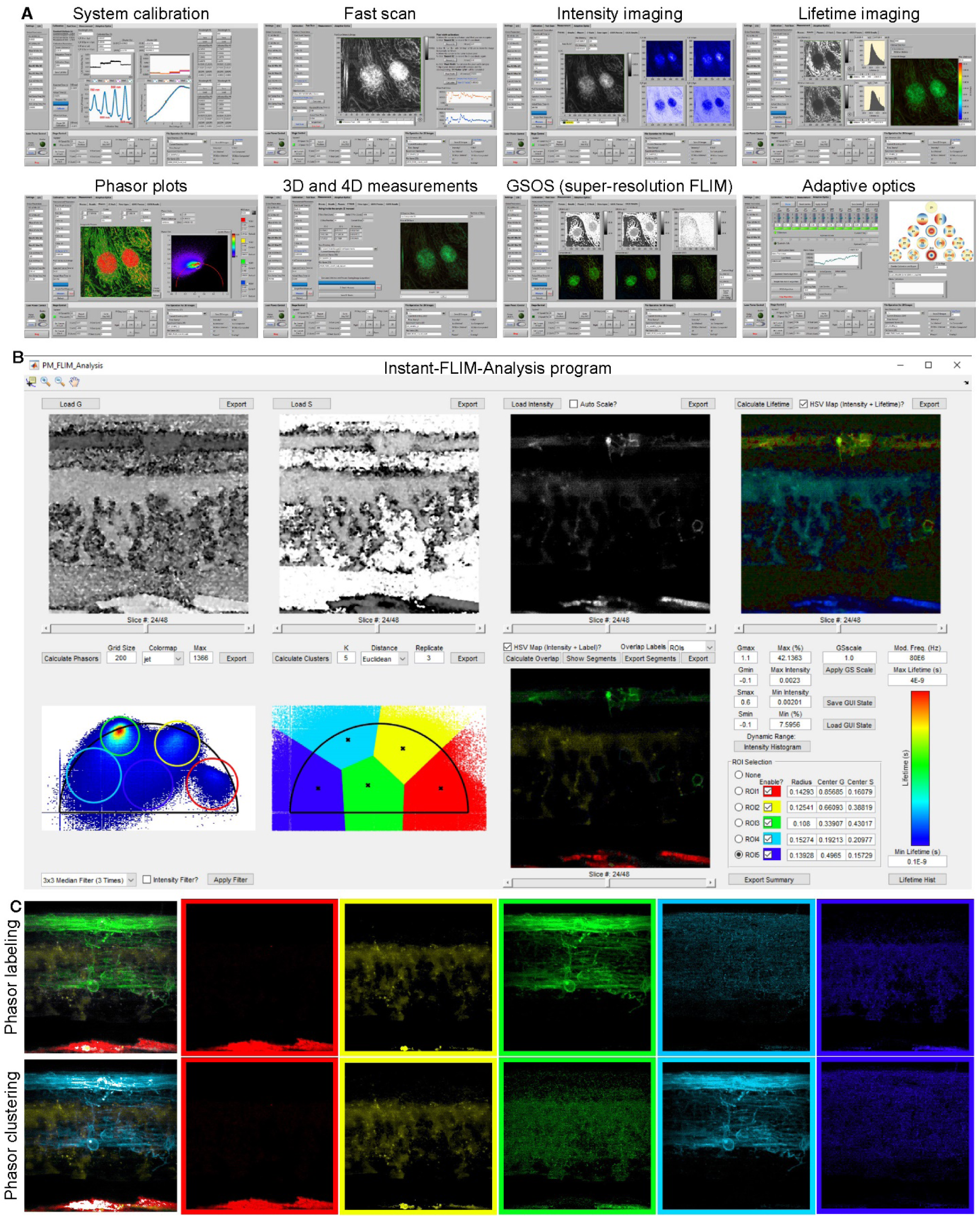
Overview of the *Instant-FLIM-Control* and *Instant-FLIM-Analysis* software. (**A**) “System calibration” module calibrates the instant FLIM system using a lifetime standard. “Fast scan” module allows fast acquisition (>1 frames per second using galvo scanners) of two-photon intensity images, which can be used to locate the focal position, to find a ROI, and to generate image-based metrics for the AO optimization algorithms. “Intensity imaging”, “Lifetime imaging”, and “Phasor plots” modules simultaneously acquire and instantaneously generate intensity images, lifetime images, and phasor plots. “3D and 4D measurements” module configures 3D and 4D instant FLIM measurements and displays the results. “GSOS (super-resolution FLIM)” module permits two-step GSOS super-resolution FLIM using two instant FLIM images of the same field-of-view captured under different excitation powers. “Adaptive optics” module optimizes the Zernike coefficients applied to the deformable mirror using the three AO optimization algorithms described in Section S4. (**B**) Screenshot of the *Instant-FLIM-Analysis* program where the 3D intensity and phasor stacks of a living *Tg(sox10:megfp)* zebrafish embryo (2 dpf) acquired with an instant FLIM system were imported and analyzed. (**C**) Maximized z-projections of the exported 3D stacks segmented by the phasor labels (top) and the phasor clusters (bottom, K=5) shown in (B).

**Fig. S6.**
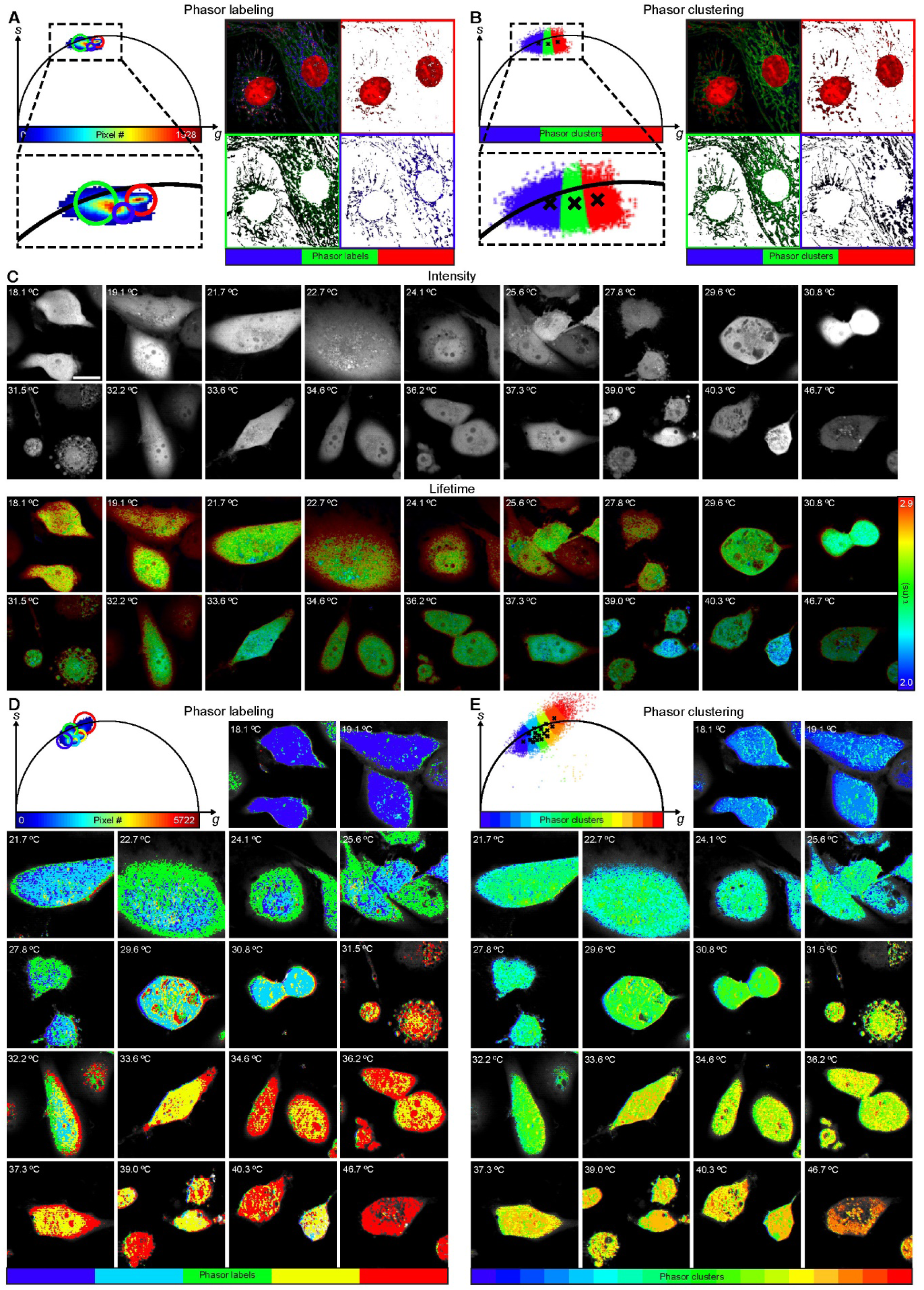
Instant FLIM with phasor labeling and phaser clustering techniques. **(A)** Demonstration of the phasor labeling technique, where ROIs are manually drawn on the phasor plot of the fixed BPAE cells to label different cellular structures shown on the right. (**B**) Demonstration of the phasor clustering technique, where the phasors of the fixed BPAE cells are automatically clustered using the K-means clustering algorithm (K=3) to denote different cellular structures shown on the right. (**C**) Two-photon fluorescence intensity and lifetime images of MDA-MB-231-EGFP cells under 18 different temperatures acquired with an instant FLIM system. (**D**) Application of the phasor labeling technique to the instant FLIM results of the MDA-MB-231-EGFP cells in (C). (**E**) Application of the phasor clustering technique (K=18) to the instant FLIM results of the MDA-MB-231-EGFP cells in (C). As temperature increases, the GFP lifetimes decrease and the phasors shift from bottom-left to top-right of the phasor plot. Scale bar, 20 μm.

**Fig. S7.**
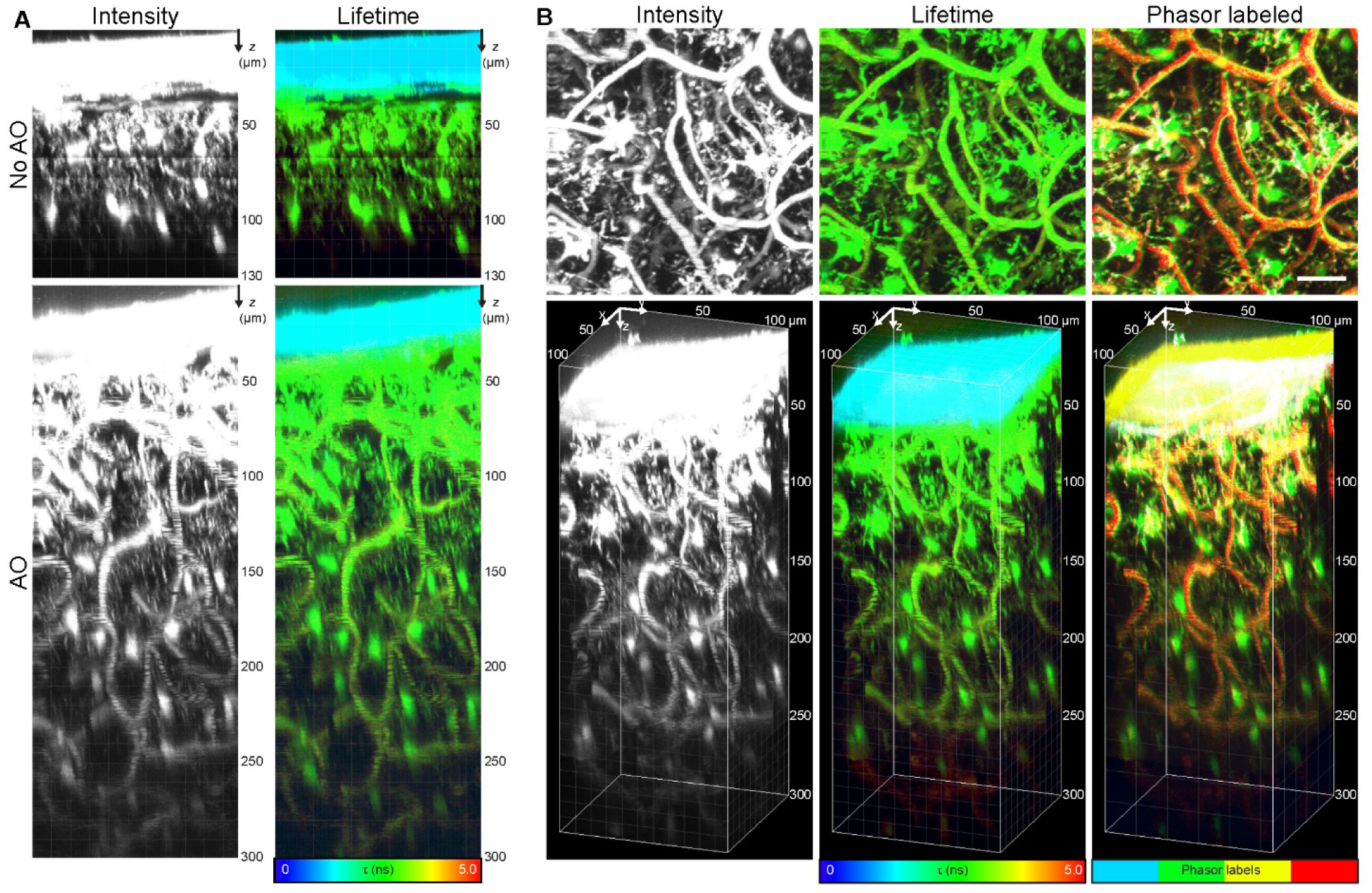
Instant FLIM with adaptive optics enabling through-skull *in vivo* lifetime imaging of intact mouse brains. (**A**) *In vivo* two-photon fluorescence intensity and lifetime maximized x-projection 3D images of an intact Cx3cr1-GFP/+ mouse brain acquired with an instant FLIM system without (top) and with (bottom) the adaptive optics (AO) module. (**B**) Skull-excluded maximized z-projections (top) and skull-included 3D reconstructions (bottom) of the *in vivo* intensity, lifetime, and phasor labeled 3D stacks of the intact mouse brain acquired with an instant FLIM system with the AO module. The microglia were labeled with EGFP, and the blood vessels were injected with Texas Red-Dextran. The excitation power was gradually increased from 8.53 mW to 21.13 mW to compensate for the signal loss as the imaging depth increased. Scale bar, 20 μm.

**Fig. S8.**
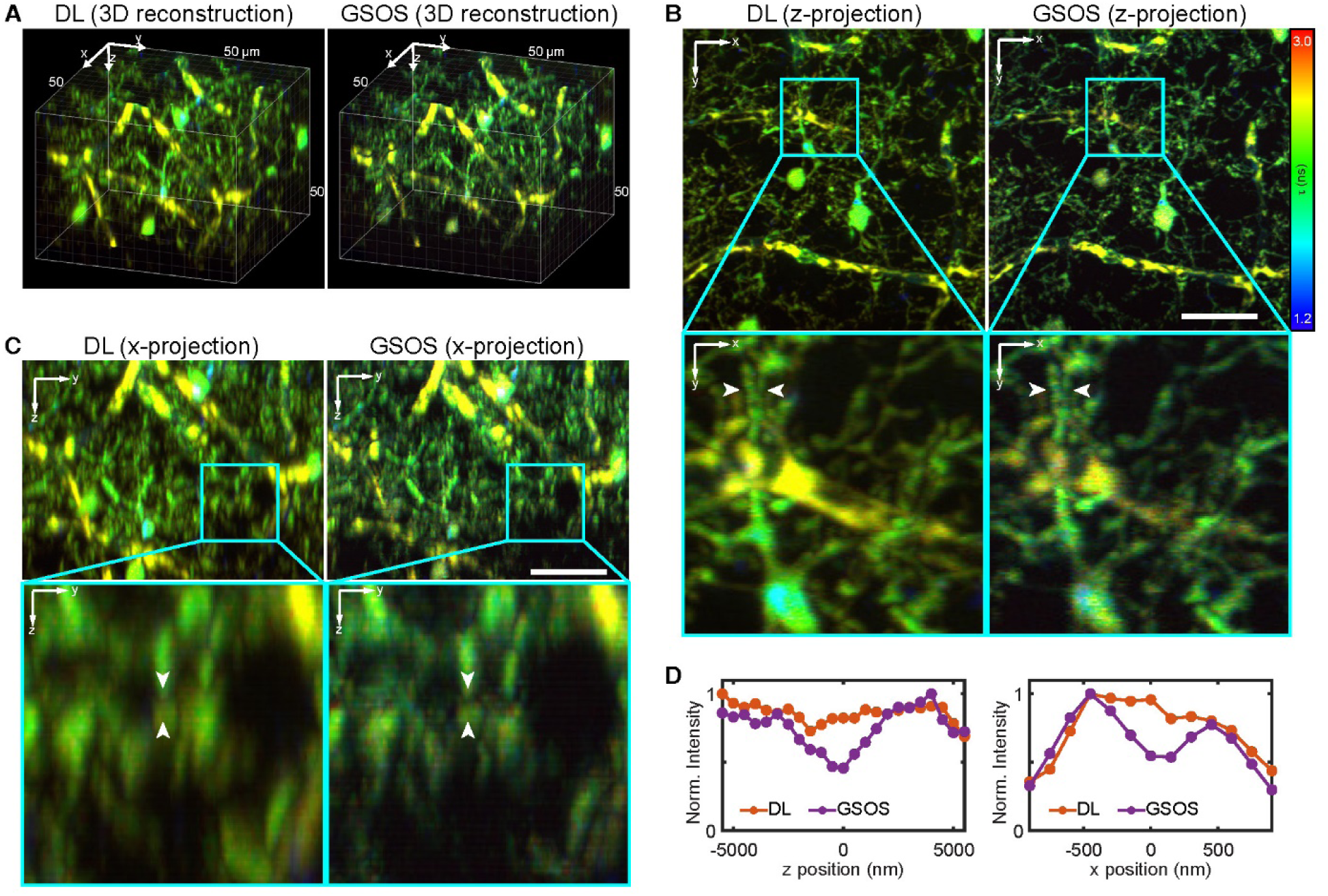
Super-resolution instant FLIM enabled by GSOS. (**A**) 3D fluorescence lifetime stacks of a fixed Cx3cr1-GFP/+ mouse brain (microglia labeled with EGFP; blood vessel injected with Dextran 594) captured by an instant FLIM system conventionally and combined with the two-step GSOS technique. The conventional diffraction-limited (DL) stack was imaged at a power of 12.31 mW, and the GSOS stack was generated by linear combining two stacks imaged at 12.31 mW and 13.44 mW, respectively. (**B**) Maximized z-projections of the 3D stacks in (A). Insets: magnified views. (**C**) Maximized x-projections of the 3D stacks in (A). Insets: magnified views. (**D**) Line profiles of the normalized intensity values at the positions of the white arrowheads in the DL and GSOS stacks in (B) and (C). Norm., normalized. Scale bars, 20 μm.

**Table S1.**
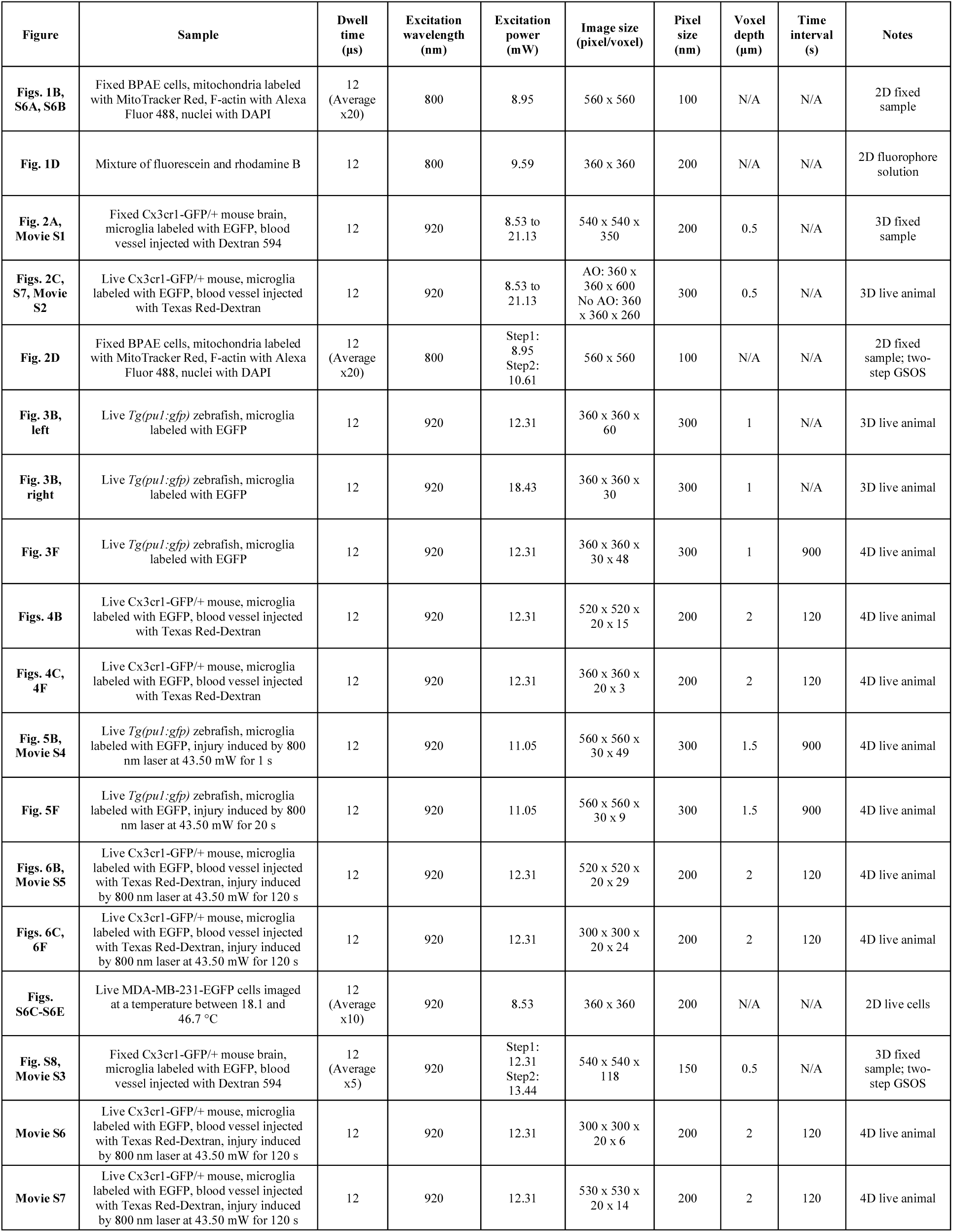
Acquisition parameters for all data. A table showing the acquisition parameters for all the data used in this work.

**Table S2.** Parts and price list for instant FLIM. A table showing the budget of all the components necessary for upgrading a conventional two-photon laser scanning microscope (including data acquisition devices) to an instant FLIM system.

| Vendor | Part number | Qty | Price | Description and link |
| --- | --- | --- | --- | --- |
| Instant FLIM components |  |  |  |  |
| ThorLabs | EF502 | 4 | \$69.86 | Low Pass Filter (LPF), DC-100 kHz, BNC Connector<br>https://www.thorlabs.com/thorproduct.cfm?partnumber=EF502 |
| Mini-Circuits | BLP-90+ | 2 | \$35.45 | Low Pass Filter (LPF), DC-81 MHz, BNC Connector<br>https://www.minicircuits.com/WebStore/dashboard.html?model=BLP-90%2B |
| Mini-Circuits | ZX60-P103LN+ | 6 | \$69.95 | Low Noise Amplifier (LNA), 50-3000 MHz, SMA Connector<br>https://www.minicircuits.com/WebStore/dashboard.html?model=ZX60-P103LN%2B |
| Mini-Circuits | ZSC-4-3+ | 2 | \$56.95 | 4-Way Power Splitter, 0.25-250 MHz, BNC Connector<br>https://www.minicircuits.com/WebStore/dashboard.html?model=ZSC-4-3%2B |
| Mini-Circuits | JSPHS-150+ | 8 | \$34.95 | Phase Shifter (PS), 100-150 MHz, Surface Mount<br>https://www.minicircuits.com/WebStore/dashboard.html?model=JSPHS-150%2B |
| Mini-Circuits | TB-152+ | 8 | \$29.95 | Board to Surface Mount Phase Shifters, SMA Connector<br>https://www.minicircuits.com/WebStore/dashboard.html?model=TB-152%2B |
| Mini-Circuits | ZAD-3H+ | 4 | \$50.95 | Level 17 Frequency Mixer, 0.05-200 MHz, BNC Connector<br>https://www.minicircuits.com/WebStore/dashboard.html?model=ZAD-3H%2B |
| Mini-Circuits | ZFBT-282-1.5A+ | 1 | \$59.95 | Bias Tee, 10-2800 MHz, SMA Connector<br>https://www.minicircuits.com/WebStore/dashboard.html?model=ZFBT-282-1.5A%2B |
| Total cost: | \$1,666.89 | | | |
| Cables and adapters |  |  |  |  |
| Mini-Circuits | 141-24BM+ | 4 | \$19.45 | 24" Coaxial Cable, BNC Connector<br>https://www.minicircuits.com/WebStore/dashboard.html?model=141-24BM%2B |
| Mini-Circuits | 141-18BM+ | 18 | \$18.45 | 18" Coaxial Cable, BNC Connector<br>https://www.minicircuits.com/WebStore/dashboard.html?model=141-18BM%2B |
| Mini-Circuits | 086-6BM+ | 6 | \$15.75 | 6" Coaxial Cable, BNC Connector<br>https://www.minicircuits.com/WebStore/dashboard.html?model=086-6BM%2B |
| Mini-Circuits | 086-2SM+ | 8 | \$10.75 | 2" Coaxial Cable, BNC Connector<br>https://www.minicircuits.com/WebStore/dashboard.html?model=086-2SM%2B |
| Mini-Circuits | SF-BF50+ | 6 | \$3.95 | SMA-F to BNC-F Adapter<br>https://www.minicircuits.com/WebStore/dashboard.html?model=SF-BF50%2B |
| Mini-Circuits | SM-BF50+ | 12 | \$3.95 | SMA-M to BNC-F Adapter<br>https://www.minicircuits.com/WebStore/dashboard.html?model=SM-BF50%2B |
| Mini-Circuits | SM-BM50+ | 6 | \$3.95 | SMA-M to BNC-M Adapter<br>https://www.minicircuits.com/WebStore/dashboard.html?model=SM-BM50%2B |
| Total cost: | \$685.20 | | | Note: cheaper cables and adapters are available from other vendors |

**Movie S1. Instant FLIM imaging of a fixed mouse brain**.

Slices and 3D reconstructions of two-photon fluorescence intensity, lifetime, and phasor labeled stacks of a fixed Cx3cr1-GFP/+ mouse brain (microglia labeled with EGFP; blood vessel injected with Dextran 594) measured with an instant FLIM system. The stacks were captured from the surface to the depth of 175 μm inside the tissue.

**Movie S2. Through-skull *in vivo* instant FLIM imaging of intact mouse brains with adaptive optics**.

Slices and 3D reconstructions of two-photon fluorescence intensity and lifetime stacks of a living Cx3cr1-GFP/+ mouse brain acquired with an instant FLIM system without (left) and with (right) an adaptive optics (AO) module. The “No AO” and “AO” stacks were captured from the surface of the skull to 130 μm and 300 μm inside the brain, respectively. The excitation power was gradually increased from 8.53 mW to 21.13 mW to compensate for the signal loss as the imaging depth increased.

**Movie S3. Super-resolution instant FLIM imaging of a fixed mouse brain**.

3D reconstructed fluorescence lifetime stacks of a fixed Cx3cr1-GFP/+ mouse brain (microglia labeled with EGFP; blood vessel injected with Dextran 594) captured by an instant FLIM system conventionally (left) and combined with the two-step GSOS technique (right). The stacks were captured from the surface to the depth of 59 μm inside the tissue. The diffraction-limited (DL) conventional stack was imaged at a power of 12.31 mW, and the GSOS stack was generated by linear combining two stacks imaged at 12.31 mW and 13.44 mW, respectively.

**Movie S4. 4D *in vivo* instant FLIM imaging in an injured zebrafish brain**.

4D reconstructions of the two-photon fluorescence intensity, lifetime, and lifetime surface rendering stacks of the microglia in a living *Tg(pu1:gfp)* zebrafish brain at 4 dpf acquired with an instant FLIM system. The 4D stacks were captured every 15 minutes and the totally imaging session was 12 hours long. A laser injury was induced at *t* = 30 minutes by an 800 nm laser at 43.50 mW lasting for 1 second. The site of the injury is denoted with dashed yellow circles. The dynamics as well as the lifetime changes of the zebrafish microglia responding to the injury can be observed with the 4D movie.

**Movie S5. 4D *in vivo* instant FLIM imaging in an injured mouse brain**.

4D reconstructions of the two-photon fluorescence intensity and lifetime stacks of the microglia in a living Cx3cr1-GFP/+ mouse brain acquired with an instant FLIM system. The 4D stacks were captured every 2 minutes and the totally imaging session took 56 minutes. A laser injury was induced at *t* = 4 minutes by an 800 nm laser at 43.50 mW lasting for 120 seconds. The site of the injury is denoted with dashed yellow circles. The dynamics and the lifetime changes of the mouse microglia responding to the injury can be seen with the 4D movie.

**Movie S6. 4D *in vivo* lifetime and phasor imaging in injured mouse brains**.

Two-photon fluorescence intensity and lifetime maximized z-projections of the 4D stacks as well as their phasor plots for each time point of the microglia in a living Cx3cr1-GFP/+ mouse brain acquired with an instant FLIM system. A laser injury was induced at *t* = 0 by an 800 nm laser with a power of 43.50 mW lasting for 120 s. The sites of the injury are denoted with dashed yellow circles. The stacks were captured at 8 different injury sites from one of two mice.

**Movie S7. Phasor labeling and phasor clustering techniques applied to 4D *in vivo* instant FLIM stacks of an injured mouse brain**.

Reconstructed 4D stacks of the microglia in a living Cx3cr1-GFP/+ mouse brain segmented with the phasor labeling (left) and phasor clustering (K=5) (right) techniques. A laser injury was induced at *t* = 0 by an 800 nm laser with a power of 43.50 mW lasting for 120 s. The site of the injury is denoted with dashed yellow circles. The cellular structures segmented by the phasor labels or clusters all responded to the injury.

